# Synaptic changes in pallidostriatal circuits observed in parkinsonian model triggers abnormal beta synchrony with accurate spatio-temporal properties across the basal ganglia

**DOI:** 10.1101/2023.03.07.531640

**Authors:** Shiva Azizpour Lindi, Nicolas P. Mallet, Arthur Leblois

## Abstract

Excessive oscillatory activity across basal ganglia (BG) nuclei in the *β* frequencies (12–30Hz) is a hallmark of Parkinson’s disease (PD). While the link between oscillations and symptoms remains debated, exaggerated *β* oscillations constitute an important biomarker for therapeutic effectiveness in PD. The neuronal mechanisms of *β-*oscillation generation however remain unknown. Many existing models rely on a central role of the subthalamic nucleus (STN) or cortical inputs to BG. Contrarily, neural recordings and optogenetic manipulations in normal and parkinsonian rats recently highlighted the central role of the external pallidum (GPe) in abnormal *β* oscillations, while showing that the integrity of STN or motor cortex is not required. Here, we evaluate the mechanisms for the generation of abnormal *β* oscillations in a BG network model where neuronal and synaptic time constants, connectivity, and firing rate distributions are strongly constrained by experimental data. Guided by a mean-field approach, we show in a spiking neural network that several BG sub-circuits can drive oscillations. Strong recurrent STN-GPe connections or collateral intra-GPe connections drive gamma oscillations (*>* 40Hz), whereas strong pallidostriatal loops drive low-*β* (10-15Hz) oscillations. We show that pathophysiological strengthening of striatal and pallidal synapses following dopamine depletion leads to the emergence of synchronized oscillatory activity in the mid-*β* range with spike-phase relationships between BG neuronal populations in-line with experiments. Furthermore, inhibition of GPe, contrary to STN, abolishes oscillations. Our modeling study uncovers the neural mechanisms underlying PD *β* oscillations and may thereby guide the future development of therapeutic strategies.

**Significance statement:** In Parkinson’s disease, neural activity in subcortical nuclei called the basal ganglia displays abnormal oscillatory synchronization that constitutes an important biomarker for therapeutic effectiveness. The neural mechanisms for the generation of these oscillations remain unknown. Here, in a theoretical neuronal network model strongly constrained by anatomical and physiological data, we show that specific circuit modifications in basal ganglia connectivity during Parkinson’s disease lead to the emergence of synchronized oscillatory activity in the network with properties that strongly agree with available experimental evidence. This and future theoretical investigations of the neural mechanisms underlying abnormal neuronal activity in Parkinson’s disease are necessary to guide the future development of therapeutic strategies to ameliorate symptoms.

## Introduction

Dysfunctions of the basal ganglia (BG)-cortical network are centrally involved in movement disorders such as Parkinson’s disease (PD). According to the classical model of BG dysfunction (DeLong, 1990; Albin R., 1989), degeneration of dopaminergic neurons in PD induces an imbalance between direct and indirect BG pathways, over-inhibiting the motor system and thereby causing hypo-kinetic motor symptoms. Accordingly, inactivation of internal pallidum (GPi) or subthalamic nucleus (STN) palliates PD symptoms (Bergman et al., 1990; Benazzouz et al., 1993), whereas activating the indirect pathway generates hypokinesia in rodents (Kravitz et al., 2010). This classical model has recently been challenged. First, the external pallidum (GPe) contains at least two cell-types with different functions (Mallet et al., 2012; Fujiyama et al., 2016; Hernández et al., 2015): the Prototypic (forming the indirect pathway) and the Arkypallidal (only projecting to striatum) neurons. Second, BG nuclei display strong modifications in firing patterns in PD, with excessive oscillatory synchrony in the *β* frequency range (12–30 Hz) (Bergman et al., 1994; Hutchison et al., 2004; Bevan et al., 2002). The link between *β* oscillations and motor symptoms remains a matter of debate; while apparently not causal to motor symptoms (Leblois et al., 2007; Degos et al., 2009; Quiroga-Varela et al., 2013), they are correlated with akinesia/rigidity levels (Kühn et al., 2006; Hammond et al., 2007; Kühn et al., 2009; Sharott et al., 2014; Neumann et al., 2016). Nevertheless, the *β* power in STN or pallidal local field potentials serves as an important biomarker for deep brain stimulation in PD patients (Little and Brown, 2020; Tinkhauser et al., 2020). Unlike mean firing rate changes, the generation mechanisms of pathological oscillations remain unknown and several competing models have been proposed (reviewed in (Pavlides et al., 2015; Rubin, 2017), but also see (McCarthy et al., 2011; Corbit et al., 2016; Adam et al., 2022; West et al., 2022)). From a theoretical perspective, any neuronal network incorporating negative feedback loops with delays can generate oscillations, with time constants and delays in the loop constraining the oscillations frequency (Ermentrout et al., 2001). The first BG pattern-generator system identified was the reciprocally-connected STN-GPe circuit (Plenz and Kital, 1999; Terman et al., 2002; Bevan et al., 2002; Nevado Holgado et al., 2010). However, many recurrent inhibitory feedback loops in the BG-cortical network could generate PD-related *β-*synchronization (Pavlides et al., 2015; Rubin, 2017) and alternative circuit generators have been proposed in the cortex (Brittain and Brown, 2014), the striatum (McCarthy et al., 2011), along the hyperdirect pathway (Leblois et al., 2006) and the recently uncovered striato-pallidal circuits (Mallet et al., 2012; Corbit et al., 2016). While these previous studies were at least partially constrained by available anatomical and physiological evidence, the heterogeneity of the biophysical parameters of the underlying model makes the direct comparison of the various generation mechanisms difficult (although see (Pavlides et al., 2015)). Recently, neural recordings combined with optogenetic manipulations in normal and parkinsonian rats highlighted the central role of the GPe in the expression of *β* oscillations, while showing that it does not require the integrity of STN or motor cortex (De la Crompe et al., 2020), ruling out several of the previously suggested mechanisms. This study also highlighted that the phase relationship between various BG neural populations is a critical hint to probe which circuit is generating the *β* oscillations. Here, we evaluate the mechanisms for generation of abnormal *β* oscillations in a BG network model where time constants, synaptic strengths, connectivity patterns, and firing rate distributions are strongly constrained by available anatomical and physiological (*in vivo* and *in vitro*) rodent data. Guided by a mean-field approach, we show in a spiking neural network that the frequency of the pathological *β-*oscillations and their phase-locking properties across BG nuclei are most likely driven by the interaction between the slow-oscillating pallidostriatal loop and the fast inhibitory loop made by recurrent inhibitory collaterals between GPe prototypical neurons.

## Materials and Methods

### Model Architecture

In this article, we focus on the BG circuits that may generate synchronized oscillatory activity. As highlighted in the introduction, oscillatory activity emerges from inhibitory feedback with delay. As the output nuclei of the BG (GPi/SNr) do not send significant projections back to other BG nuclei, they do not participate in recurrent feedback circuits. Similarly, D1-expressing medium spiny neurons (D1-MSNs) in the striatum mainly participate in purely feedforward direct pathways and are unlikely to be included in a significant recurrent circuit inside the BG (but see (McCarthy et al., 2011)). In addition, their activity is strongly reduced after dopamine depletion (Mallet et al., 2006; Parker et al., 2018). For all these reasons, we do not include output nuclei and D1-MSNs in our model. We include all other major populations of the basal ganglia network. In particular, our model reflects on the recently discovered cellular dichotomy between the principal GPe cell population, the prototypic neurons (projecting to STN and (GPi/SNRr), and the arkypallidal neurons (only projecting to the striatum (Mallet et al., 2012)). Our model consists of five subpopulations, four inhibitory and one excitatory: fast-spiking interneurons (FSIs) and D2-expressing medium spiny neurons (D2) in the striatum, prototypical (Proto) and arkypallidal (Arky) neurons in the GPe, and lastly, STN excitatory neurons (Fig. 1). Henceforth when referring to these neuronal populations, we simply use Proto, Arky, STN, FSI, and D2.

**Figure 1.**
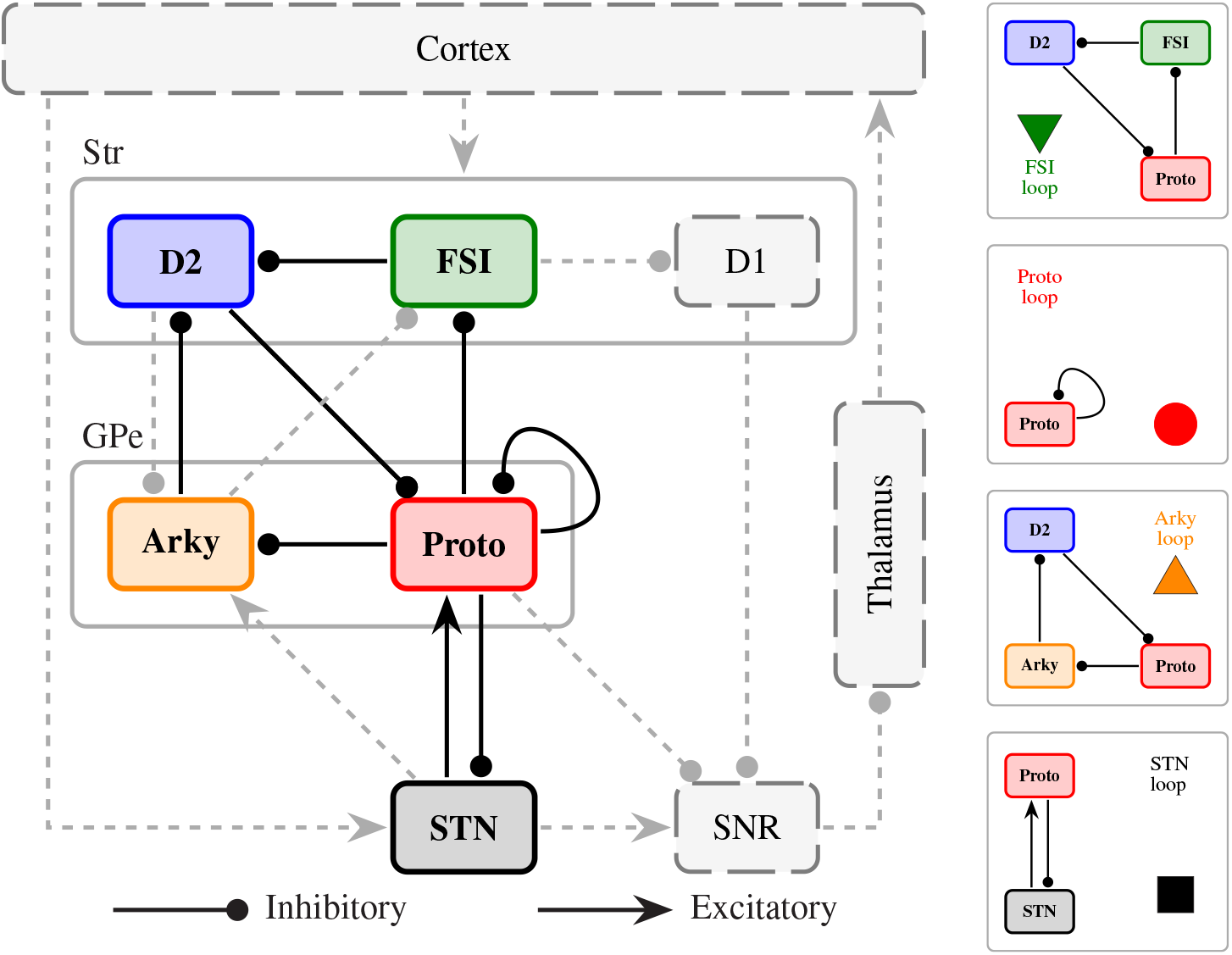
The model architecture. The model includes Prototypical (red) and Arkypallidal (orange) populations of the external globus pallidus (GPe), D2-expressing medium spiny neurons (D2, blue), and fast-spiking interneurons (FSI, green) within the striatum, and finally the subthalamic nucleus (STN, black). Arrows/dots indicate excitatory/inhibitory connections, respectively. Grey regions and connections are not included in the model. These populations make up 4 different closed negative feedback loops represented on the right: FSI loop (green triangle), Proto loop (red circle), Arky loop (orange triangle), and STN loop (black square).

We include all major connections that connect the considered neuronal populations. They contribute to several closed negative feedback loops and have the Proto subpopulation as their common node. There are four such negative feedback loops. The STN-Proto recurrent network is one which involves Proto and STN, and comprises two synaptic connections, STN to Proto (Kita and Kitai, 1991; Magill et al., 2000), and Proto to STN (Kita et al., 1983; Kita and Kita, 1994; Baufreton et al., 2009) (referred to as STN-loop in the figures). Second, there is lateral inhibition within the Proto subpopulation (Ketzef and Silberberg, 2020; Sims et al., 2008) (referred to as the Proto-loop). The two remaining are pallidostriatal loops that both include the projection from D2-MSNs to Proto neurons (Kita, 1994; Cooper and Stanford, 2000; Ketzef and Silberberg, 2020; Aristieta et al., 2021). The FSI loop has the additional connections of Proto to FSI (Bevan, 1998; Saunders et al., 2016; Mallet et al., 2012; Corbit et al., 2016) as well as FSI to D2 (Mallet et al., 2005). Lastly, the Arky-loop is the one that connects Proto to Arky (Ketzef and Silberberg, 2020; Aristieta et al., 2021), Arky to D2 (Mallet et al., 2012; Fujiyama et al., 2016) and as previously mentioned, D2 to Proto. We have deliberately ignored connections that have been shown to be significantly weaker (or not well characterized), namely the connection from STN to Arky (weaker than STN to Proto, see (Ketzef and Silberberg, 2020; Aristieta et al., 2021)), from Arky to FSI (as this projection is not well characterized and likely weaker than the selective and potent inhibition from Proto to FSI (Bevan, 1998; Saunders et al., 2016)), and the projection from D2 to Arky (weaker than D2 to Proto, see (Ketzef and Silberberg, 2020)). As explained below, all parameters of the model are extracted or strongly constrained by experimental data collected in rodents. When available, we use experimental data from rats, otherwise, we use data from mice instead and clearly indicate it in the corresponding tables.

### Population Size and Connectivity

The number of neurons in all populations of the model is scaled down compared to the real neuron population size to make the simulation computationally tractable. We consider a constant number of neurons *N*^*sim*^ when simulating all *α* sub-populations (*N*^*sim*^ = 1000 in the spiking model and *N*^*sim*^ = 100 in the rate model, see below) and follow the approach proposed by (Golomb and Hansel, 2000) to set the number of connections population size accordingly. As such, the number 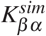 of synapses received by one neuron in population *β* from all neurons in population *α* is determined by:

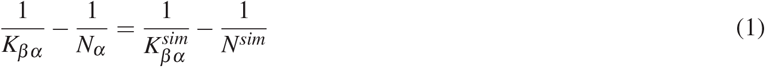

where *K*_*βα*_ is the number of connections from population *α* to a neuron in population *β* and *N*_*α*_ the size of population *α* found in the experimental literature (see Table. S1-1). This scaling ensures that the dynamical properties of the original circuit are preserved in the scaled version. The values of *K*_*βα*_ derived for each population for the spiking neuronal network with *N*^*sim*^ = 1000 are reported in Table. 1. *K*_*βα*_ are calculated similarly for *N*^*sim*^ = 100 in the rate model (values not shown here). Note that henceforth we use the notation *α*-*β* to refer to the synaptic projection from population *α* to population *β*.

**Table 1.** Number of presynaptic neurons converging onto a postsynaptic neuron.

| Parameter | Value | Reference |
| --- | --- | --- |
| $K_{Proto-D2}$ | 3100 | (Koshimizu et al., 2013) |
| $K_{STN-Proto}$ | 442 | (Baufreton et al., 2009) |
| $K_{Proto-STN}$ | 63 | (Koshimizu et al., 2013) |
| $K_{Proto-Proto}$ | 104 | (Sadek et al., 2006) |
| $K_{Arky-Proto}$ | 104 | (Sadek et al., 2006) |
| $K_{D2-FSI}$ | 51 | (Guzmán et al., 2003; Gittis et al., 2011) |
| $K_{FSI-Proto}$ | 67 | (Bevan, 1998) |
| $K_{D2-Arky}$ | 10 | Estimate (see Appendix. 2) |

$K_{\beta\alpha}$ represents the number of presynaptic neurons in population $\alpha$ converging onto one postsynaptic neuron in $\beta$ for all connections in the spiking neuronal BG network. The references on which the estimates are based are reported in the last column. See Appendix. 2 for more details on the derivation of $K_{\beta\alpha}$ from experimental observation.

### Neuronal model

While most results shown here are computed in a spiking neuronal network model, we will also consider a simplified version of the model in which neurons are represented by their firing rate (Wilson and Cowan, 1972). This model has the advantage that several aspects of its dynamics can be investigated analytically. The knowledge of the properties of this reduced model guides us in the investigation of the detailed spiking model.

#### Rate model

In order to simulate the firing activity of each neuron, we use a model similar to Leblois et al. (2006), where the dynamics of the activity of a single neuron is characterized by the rate model (Wilson and Cowan, 1972). The spontaneous firing rate of a neuron is given by 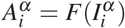, where *α* denotes the population and can be any of Proto, STN, Arky, D2, or FSI. 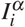 is the total input to neuron *i* where *i* = 1, ..*N*_*α*_ and *F* is its non-linear input-output transfer function. Here for simplicity, the transfer function is considered to be a piecewise linear function *F* = [*I*_*i*_ *− θ*], where [*I*_*i*_]_+_ = *I*_*i*_ for *x >* 0 and 0 otherwise with the activity threshold *θ* = 0.1. Each neuron *i* in population *α* has a synaptic contribution of 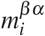 to each neuron in postsynaptic population *β*. 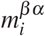 is a low-pass filtered version of the instantaneous level of activity 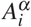 (Shriki et al., 2003), and its dynamics are governed by

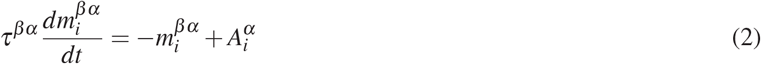

where *τ*^*βα*^ is the synaptic decay time constant of the projection from population *α* to population *β*. The simulation of the dynamics is integrated using a standard first-order forward Euler method with a time step *dt* = 0.1 ms to ensure maximum precision. Each neuron receives multiple synaptic inputs from the neurons their presynaptic populations determined by the connection matrix *J*^*βα*^. Therefore, we need to sum the inputs from all presynaptic neurons to get the net synaptic input to each postsynaptic neuron,

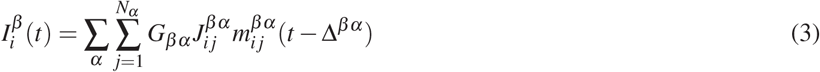

where ∆^*βα*^ is the axonal transmission delay of the projection from population *α* to population *β* (see first column in Table. 5). We simulate each population with size *N* = 100. We have also performed the simulations with size *N* = 1000 to ensure that the dynamics remain unchanged in our smaller network. Moreover, we have investigated this reduced model analytically in the case of all-to-all connectivity in the network. In this case, the steady states of the network in response to constant inputs can be obtained by solving the fixed-point equations for the dynamics (see Appendix. 1). In these states, the activities are determined by a set of linear equations that can be solved straightforwardly. The conditions for the stability of steady states can then be derived. Indeed, a fixed-point solution of the dynamical equations is stable if any small perturbation around it eventually decays over a large time. If certain perturbations increase with time, the fixed point is unstable. To investigate the stability of a steady state, we study the equations of the dynamics linearized around that state (Strogatz, 2018) (see derivation in Appendix. 1). In particular, the derivation leads to the estimation of the frequencies of the oscillatory instabilities for individual negative feedback loops in the BG network (see Appendix. 1 for more detail).

#### Spiking Neural Network (SNN)

The neurons in the SNN are simulated using the leaky-integrate and fire (LIF) model (Lapicque, 1907). In this model, the dynamics of the membrane potential of each neuron are given by

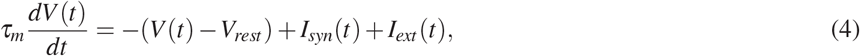

where *V* (*t*) is the membrane potential of each neuron at time *t, V*_*rest*_ the resting membrane potential (RMP), *τ*_*m*_ the membrane time constant, *I*_*syn*_(*t*) the total synaptic input, and *I*_*ext*_(*t*) the external input of each individual neuron. Once the membrane potential of the neuron reaches the action potential threshold (APth), a spike is discharged and *V* (*t*) is manually reset to the resting membrane potential (see Neuronal model for more detail on the modified *V*_*rest*_ value). The values for the RMP, APth, and *τ*_*m*_ are drawn from Gaussian distributions with respective parameters extracted from the experimental literature and summarized in Table. 2, and Table. 3. The Gaussian distribution for the RMP and *τ*_*m*_ are truncated to avoid problems with the numerical integration.

**Table 2.**
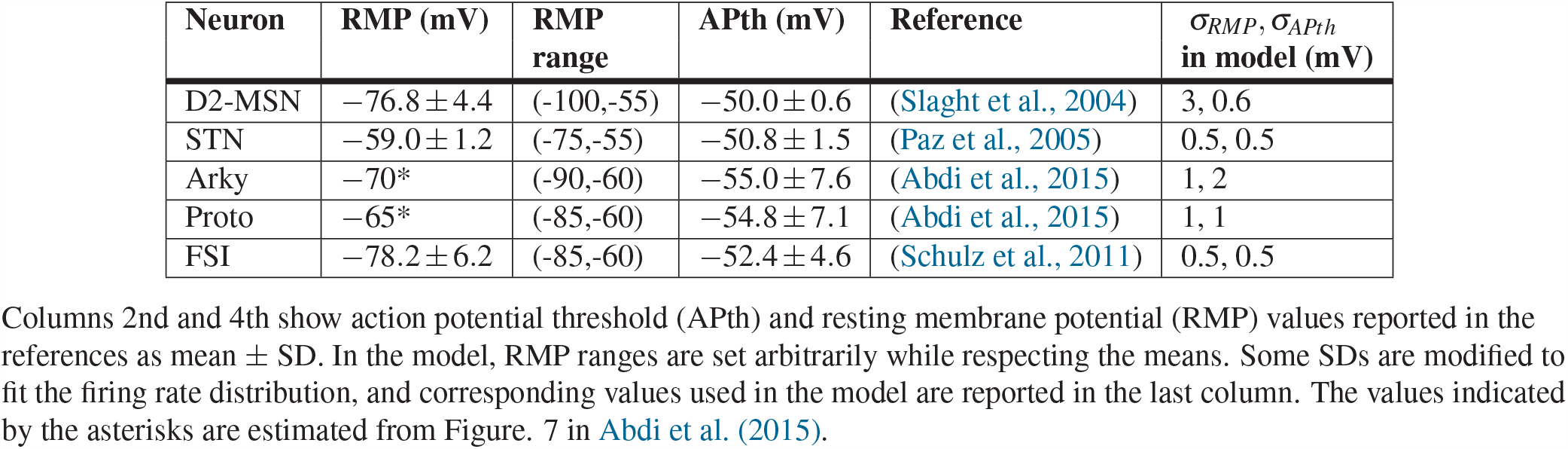
Electrophysiological properties of the action potentials.

| Neuron | RMP (mV) | RMP range | APth (mV) | Reference | $\sigma_{RMP}, \sigma_{APth}$ in model (mV) |
| --- | --- | --- | --- | --- | --- |
| D2-MSN | $-76.8 \pm 4.4$ | (-100,-55) | $-50.0 \pm 0.6$ | (Slaght et al., 2004) | 3, 0.6 |
| STN | $-59.0 \pm 1.2$ | (-75,-55) | $-50.8 \pm 1.5$ | (Paz et al., 2005) | 0.5, 0.5 |
| Arky | $-70^*$ | (-90,-60) | $-55.0 \pm 7.6$ | (Abdi et al., 2015) | 1, 2 |
| Proto | $-65^*$ | (-85,-60) | $-54.8 \pm 7.1$ | (Abdi et al., 2015) | 1, 1 |
| FSI | $-78.2 \pm 6.2$ | (-85,-60) | $-52.4 \pm 4.6$ | (Schulz et al., 2011) | 0.5, 0.5 |

**Table 3.** Membrane time constants used in the model.

| Neuron | $\tau_m$(ms) | Reference | $\sigma_{\tau_m}$ & range in model (ms) |
| --- | --- | --- | --- |
| D2-MSN | $4.9 \pm 1.6$ | (Slaght et al., 2004) | 0.5 - (2, 12) |
| STN | $5.1 \pm 2.4$ | (Paz et al., 2005) | 0.6 - (2, 10) |
| Arky | $19.9 \pm 13.0$ | (Cooper and Stanford, 2000) | 3 - (2, 100) |
| Proto | $12.9 \pm 10.2$ | (Karube et al., 2019) | 1.3 - (2.25, 45) |
| FSI | $3.1 \pm 1.1$ | (Schulz et al., 2011) | 0.3 - (1, 6) |

2nd column indicates the membrane time constants as reported in the references (mean ± SD). Some SDs are modified in the model to fit the firing rate distribution and the corresponding values used in the model are reported in the last column.

Synaptic inputs are modeled by a double exponential with independent rise and decay time constants (Fourcaud and Brunel, 2002). The sum of synaptic inputs from all presynaptic neurons *I*_*syn*_(*t*) to each neuron is given by

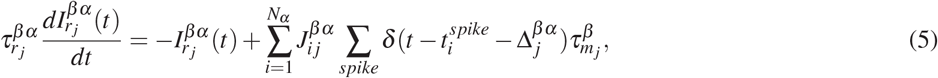

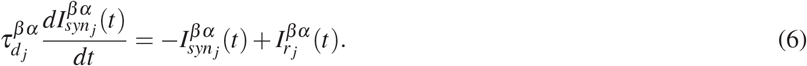

where 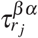 and 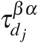 are the rise and decay time constants of the synaptic input to neuron *j* in postsynaptic population *β*. 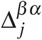 is the synaptic transmission delay from populations *α* to neuron *j* in population *β*. 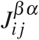 is the connectivity matrix between populations *α* and *β* where each nonzero element is equal to the synaptic weight of the corresponding synapse. 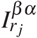 is the external input to neuron *j* in postsynaptic population *β*. As indicated by the superscripts, all of the aforementioned values are specific to each projection *β-α*. For the sake of simplicity, the time constants used to describe the incoming projections from presynaptic population *alpha* converging to neuron *j* in population *β* are all identical. However, the values of 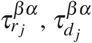 and 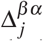 are drawn randomly from a Gaussian distribution for each postsynaptic neuron *j* = 1, …, *N*_*β*_, with means and standard deviation of the underlying distributions extracted from the experimental literature and reported in Table. 5. The connection weights 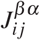 are either zero (with a probability 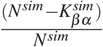) or drawn from a log-normal distribution with a standard deviation that spans one order of magnitude (Buzsáki and Mizuseki, 2014). The means of the synaptic weight distributions for the STN-Proto, Proto-STN, and Arky-Proto, are set so that the network activity resembles those of the *β-*induction experiments (see Results and Fig. 6 for more detail). Due to the lack of such experimental data for the rest of the projections, we set their weights in a way to mimic the physiological healthy states with asynchronous activity across the BG neuronal populations. The mean of the synaptic weight distributions for each projection in the healthy state is reported in Table. 4.

**Table 4.**
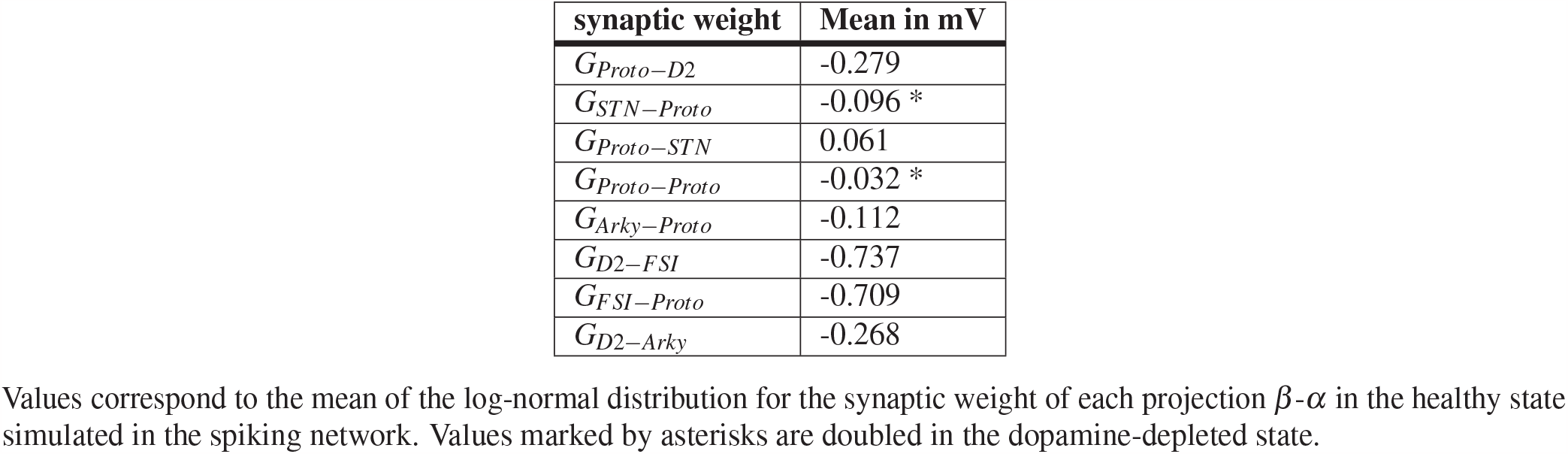
Mean of the synaptic weight distributions.

| synaptic weight | Mean in mV |
| --- | --- |
| $G_{Proto-D2}$ | -0.279 |
| $G_{STN-Proto}$ | -0.096 * |
| $G_{Proto-STN}$ | 0.061 |
| $G_{Proto-Proto}$ | -0.032 * |
| $G_{Arky-Proto}$ | -0.112 |
| $G_{D2-FSI}$ | -0.737 |
| $G_{FSI-Proto}$ | -0.709 |
| $G_{D2-Arky}$ | -0.268 |

Values correspond to the mean of the log-normal distribution for the synaptic weight of each projection $\beta$ - $\alpha$ in the healthy state simulated in the spiking network. Values marked by asterisks are doubled in the dopamine-depleted state.

**Table 5.**
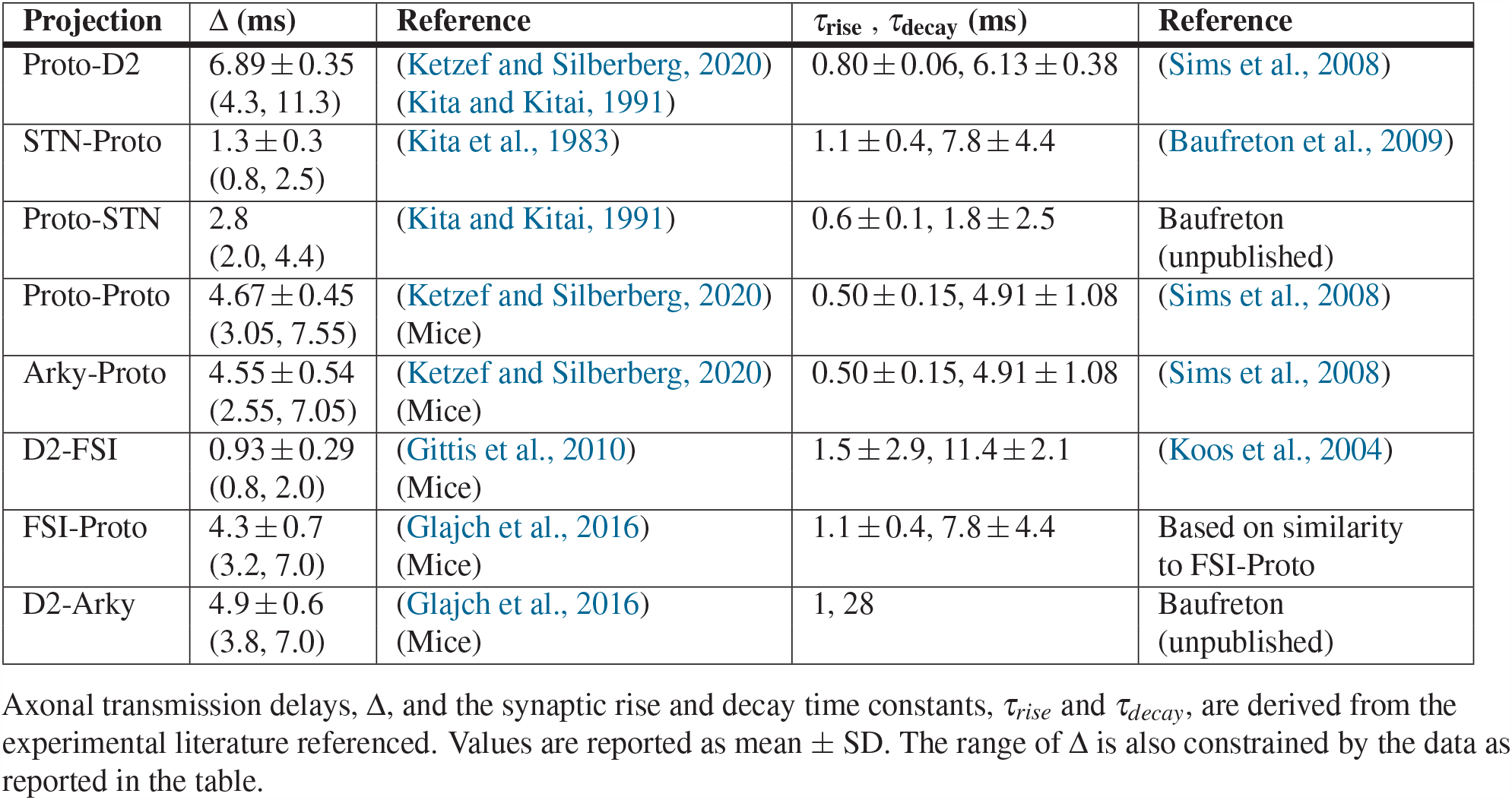
Electrophysiological time constants for each synaptic connection.

The external input given to the neurons in each population is also drawn from a Gaussian distribution. The mean I-F (Input-Firing rate) curve of the population is empirically derived for a set of external inputs with varying means and a constant standard deviation. Then, based on whether the desired firing rate is low (where the I-F curve is non-linear) or high (linear section of the curve), either a sigmoid or a line is fitted to the simulation data points. Subsequently, the mean input required to obtain the firing rate reported in the experiments for each population is extrapolated from its respective fitted curve. The standard deviation of the external input is tuned to replicate the firing rate distributions reported in the experimental literature (Table. 6). Wherever we were unable to replicate the firing rate distribution solely by manipulating the standard deviation of the external input, we tried tuning it by changing the standard deviations of *τ*_*m*_ and/or RMP, and/or APth. The standard deviations of these parameters are reported in Table. 2 and Table. 3.

#### Membrane potential initialization

Spurious synchrony can be present in the network due to the initialization of the membrane potentials of neurons (setting *V* (*t* = 0)). Such spurious synchrony would be mirrored in other synaptically connected populations and interfere with the network oscillations which we are interested to look at. We, therefore, run a 2-second long simulation to let the network reach its stable dynamical state. We then compute the distribution of membrane potential values for each neuron throughout that simulation and initialize all subsequent simulations with the corresponding membrane potential distributions.

#### Numerical integration

We use the second-order Runge-Kutta (RK2) method to numerically integrate the membrane potential at each time step. However, the discontinuity of the LIF model at the spike times can cause numerical errors. So the time steps over which the equation is integrated should be kept small in order to keep the errors to a minimum. This can slow down the simulation dramatically. As a solution, we use a linear interpolation method proposed by Hansel et. al (Hansel et al., 1998) when updating the membrane potential at spike times. The interpolated value of the membrane potential at the time step after the spike time, *V* (*t* + ∆*t*), is given by

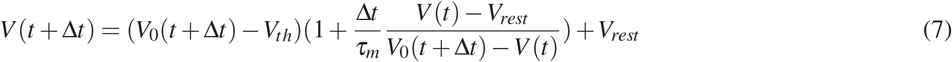

Where *V*_0_(*t* + ∆*t*) is calculated by the standard RK2 at the spike time and *V*_*th*_ is the action potential threshold. In this RK2 method with linear interpolations, the potential is reset to the value given by the Eq. 7 instead of *V*_*rest*_. When using the RK2 integration with the aforementioned interpolation, a time step of 0.25 ms results in a 10% error in coherence as compared to the exact analytical solution, whereas 0.063 results in a 1% error (Hansel et al., 1998). With this information, we chose to use a time step of 0.1 ms in our simulations. We also performed sample simulations with a 0.05 ms times step and did not observe any significant change in the resulting dynamics.

### Data Analysis

#### Population firing rate

The firing rate of all the neurons in each population is averaged in each time bin to yield the average population firing rate as shown in the figures. In the SNN, the average population firing rate is further smoothed using a moving average window of 5 ms.

#### Power spectrum

The power spectrum of the average firing rate activities is derived using the Welch method in the *scipy*.*signal* library (Welch, 1967) using 50% overlapping 1-second long windows. The simulations are run for 20.3 and 25.3 seconds for the rate model and the spiking neural network (SNN), respectively. The first 300 ms is discarded to discard the transient dynamics that depend on initialization. In addition, the SNN simulations are repeated 3 times each with different network initializations to yield a mean and standard error of the mean (SEM) for each frequency.

#### Phase histogram

The *β-*filtered mean population firing rate of the Proto neurons is used as the reference signal for the calculation of the linear phase histograms. The phase of each spike is measured relative to the two closest peaks of the reference signal in time, one before and the other after. Then, the linear phase histogram of the spikes from each neuron is calculated. Finally, the phase histogram of all neurons is subjected to Rayleigh’s test and only those that pass are taken into account. The value at which the phase histogram is maximized for each neuron is used to create the box plots in the phase histogram plots.

#### Cross-correlation

The spike cross-correlations (CC) were calculated using the *scipy*.*signal* library. 2000 random pairs of neurons are chosen for each *β-α* connection without pair replacement. Then, the CC for each pair is individually calculated using the 25-second long spike trains with 0.1 ms bin size time resolution. Subsequently, the results of coherence for all the pairs are reported as mean ± SEM. The CCs were sampled for every 2.5 ms of lag and plotted across the [− 0.2, +0.2] second lag window in order to compress the data presented in the plots.

#### Coherence

The coherence is measured using the *scipy*.*signal* library. 1000 random pairs of neurons are chosen for each *β-α* connection without pair replacement. Then, the coherence for each pair is individually calculated using the 25-second long spike trains with 0.1 ms bin size time resolution. The results of coherence for all the pairs are reported as mean ± SEM for every frequency bin.

#### Stable oscillation

We introduce a metric to help us detect the transition boundary between the steady state and the stable oscillatory regime. This metric is defined as the ratio of the peak amplitudes of the firing activity in the last cycle relative to the second cycle. Once this metric reaches the value of 1.01 and is still lower than 1.2, it is considered that the network has transitioned to a stable oscillatory regime.

## Code availability

The simulation of the network model and the data analysis are all performed using custom code in Python available at https://github.com/Shiva-A-Lindi/BG_Oscillations.

## Results

Building on the abundant anatomical and physiological data from rodents in normal and parkinsonian conditions and the well-established abnormal *β-*band synchronization present in the BG of parkinsonian rats, we study how various negative feedback loops inside a model network of rat BG circuits may generate abnormal *β-*band oscillations. We follow a two-step approach (see Materials and Methods for the details). We first rely on a simplified model where neuronal activity is represented by its firing rate and a limited number of parameters determine the frequency of oscillations in each closed-loop circuit. Guided by the analysis of this reduced model, we then analyze the properties of a spiking neuronal network model with the same architecture where each neuron is represented by its membrane potential. This realistic model allows us to faithfully represent the firing properties of BG neurons and is a more accurate representation of the circuit dynamics underlying normal and pathological BG activity. Recent evidence shows that the GPe is a necessary hub for the generation of the *β* oscillations in parkinsonian rats (De la Crompe et al., 2020), and our model, therefore, includes all closed loop BG circuits that include at least one GPe neuronal population. The GPe consists of two distinct subpopulations, the Prototypical (Proto) and the Arkypallidal (Arky) neurons (Mallet et al., 2012). The Prototypical neurons send projections to the striatum, contacting mainly the fast-spiking interneurons (FSIs) (Bevan, 1998; Saunders et al., 2016; Mallet et al., 2012; Corbit et al., 2016). This connection together with projections from FSIs to D2-MSNs, and D2-MSNs to back to Proto neurons, creates a candidate feedback loop (Corbit et al., 2016), the FSI-loop. Similarly, Arky neurons send projections to the striatum and contact D2-MSNs (Mallet et al., 2012; Fujiyama et al., 2016). Therefore, considering the lateral inhibition from Proto to Arky creates another pallidostriatal loop, the Arky-loop. The well-established recurrent network between the STN and GPe (Proto) (Plenz and Kital, 1999; Terman et al., 2002; Nevado Holgado et al., 2010; Kumar et al., 2011; Pasillas-Lépine, 2013; Nevado-Holgado et al., 2014; Pavlides et al., 2015; Shouno et al., 2017; Koelman and Lowery, 2019; Chen et al., 2020) and the recurrent lateral inhibition between Proto neurons (Ketzef and Silberberg, 2020; Aristieta et al., 2021) also constitute negative feedback loop circuits. All the loops and connections are represented in Fig. 1.

### Wide range of frequencies from BG oscillation-generator circuits

As mentioned in the introduction, a neuronal circuit forming a closed negative feedback loop can generate oscillations if the strength of its feedback is sufficient (Ermentrout et al., 2001). There are four different negative feedback loops in our BG network model that can generate oscillations, as detailed above (FSI-D2-Proto loop, D2-Proto-Arky loop, STN-Proto loop, and Proto recurrent loop, see Fig. 1). Each loop can be considered as a dynamical system with a steady state (or fixed point) in which all populations have a constant activity level, which is stable if any small perturbation around it eventually decays over time, unstable otherwise. The stability of a steady state and the nature of the instabilities can be studied by solving the equations of the dynamics linearized around that state (Strogatz, 2018). In the case of negative feedback loops, an oscillatory instability arises for sufficiently large overall gain, defined as the product of the synaptic weights along the connections of the loop (see Appendix. 1 for the detailed calculation). The frequency of the oscillatory instability in each loop is a function of the axonal delays and synaptic time constants of the various synaptic connections of the loop (see Appendix. 1). We compare the theoretically derived frequency of the oscillatory instability with the actual frequency of oscillations observed in simulations of each loop when it is isolated from the complete network in the rate model (see Materials and Methods). Synaptic weights of all connections in the loop circuit are set as to reach sufficient feedback to drive stable oscillations and measure the frequency of oscillations in the firing rate time course (see Materials and Methods for the definition of stable oscillation). The synaptic time scales of inhibition are widely debated, especially in the studies supporting the STN-GPe as the generator of *β* oscillations with values ranging from 7 to 20 ms. (Plenz and Kital, 1999; Terman et al., 2002; Nevado Holgado et al., 2010; Kumar et al., 2011; Pasillas-Lépine, 2013; Nevado-Holgado et al., 2014; Pavlides et al., 2015; Shouno et al., 2017; Koelman and Lowery, 2019; Chen et al., 2020). Therefore, we fix the decay time constant for the excitatory projection from STN to Proto to *τ*_*Proto−STN*_ = 6 ms and vary the decay time constant of inhibitory synapses *τ*_*inh*_ for all the inhibitory projections in the model in the range of 4 to 24 ms. For all values of *τ*_*inh*_, we compare the theoretically-derived frequency of the oscillatory instability with the frequency of oscillations observed in rate model simulations (Fig. 2A). The four considered loop circuits induce oscillations in a wide range of frequencies ranging from 5 to 80Hz. Indeed, the oscillation frequencies derived from simulations are well predicted by our theoretical derivation. Importantly, across the wide range of inhibitory decay time constants considered here, the STN-GPe network does not produce *β-*band oscillations but rather oscillates in the *γ* range (35-50Hz). Similarly, the short feedback provided by recurrent connections between Proto neurons drives *γ*-band oscillations with even higher frequencies (60-70 Hz). On the other hand, both pallidostriatal loops generate frequencies close to the lower boundary of the *β-*band (5-25 Hz, Fig. 2A).

**Figure 2.**
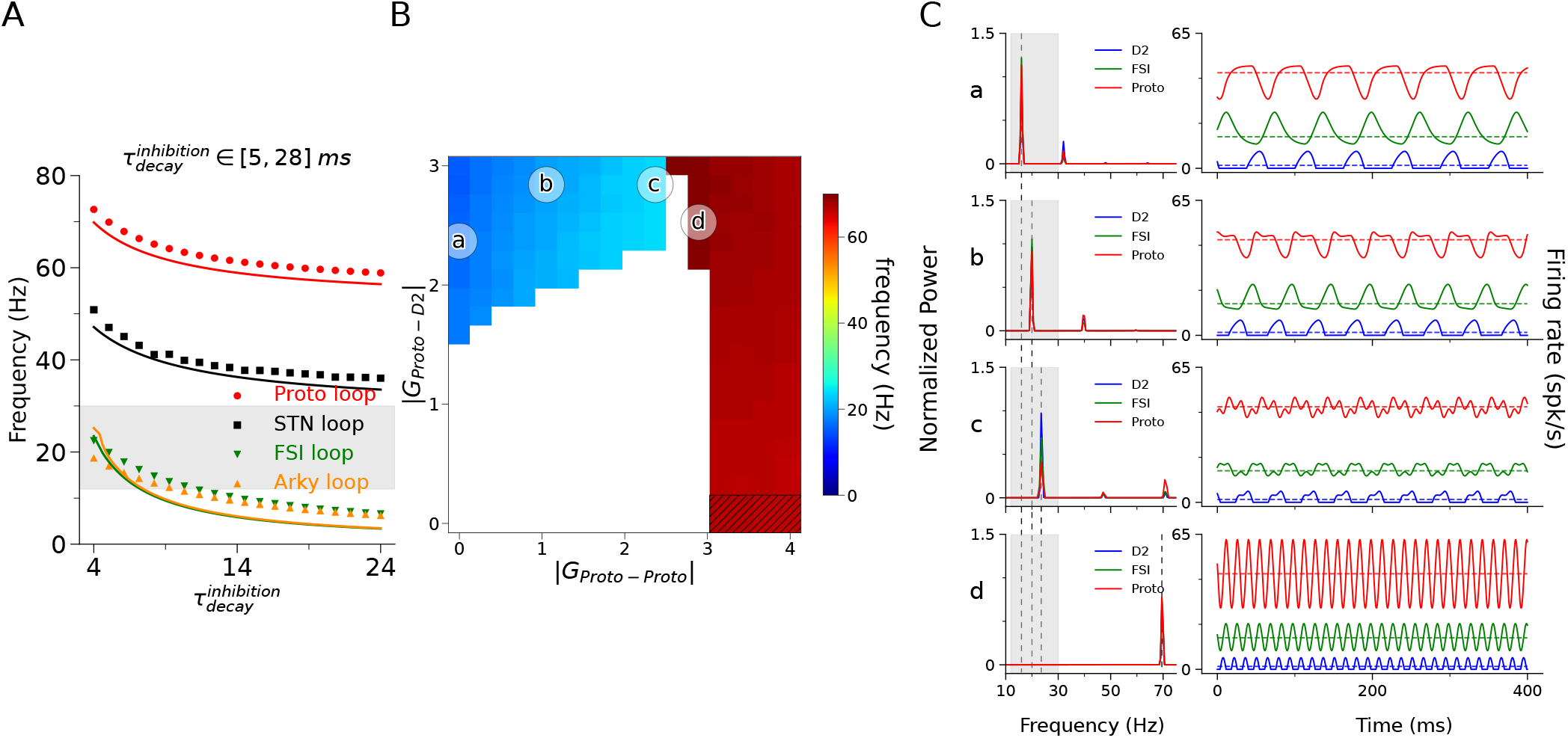
Pallidostriatal loop with the help of lateral Proto-Proto inhibition can generate *β-*band frequency oscillations in rate model. A) Frequency of stable oscillations as a function of inhibitory synaptic decay time constant *τ*_*inhibition*_ for each individual loop. The shaded area indicates the *β* band (12-30Hz). The scatter plot shows results from the rate model simulation whereas solid lines are estimates from theory (for more details see Appendix. 1). The range of values reported in experimental studies for the inhibitory synaptic time scale is shown above the plot. B) Phase space of joint FSI- and Proto-loops. All the synaptic weights in the FSI-loop increase along the y-axis though only the values of *G*_*Proto − D*2_ are indicated (*G*_*Proto − D*2_ = *G*_*D*2 *− FSI*_ = *G*_*FSI − Proto*_). The color of each point in the phase space represents the frequency of stable oscillations corresponding to the network with those weights. Whereas, the white indicates that the network is in steady state. Lastly, the hatched area indicates where only the Proto population has reached the stable oscillatory regime. C) The normalized power spectrum (left) and mean firing activity of each population for the highlighted points in (B) (right). All data points represent a 20-second simulation of 100 neurons in each population with 0.1 ms time bins.

### Interactions between multiple BG circuits including the GPe but not the STN can lead to the emergence of *β*-like activity

In the full BG network, oscillations may be generated as a result of interactions between two or more loops. Since STN is not necessary for the induction of experimentally observed *β-*band oscillations in parkinsonian rats (De la Crompe et al., 2020), the pallidostriatal loops (FSI-D2-Proto loop and D2-Proto-Arky loop) may interact with the recurrent Proto loop to drive pathological oscillations in PD. Interestingly, the synaptic connections between FSI and D2 striatal neurons and the recurrent connections between Proto neurons are strongly affected by dopamine depletion (Gittis et al., 2011; Miguelez et al., 2012), and see below for more details). Therefore, we investigate the possible interaction between the FSI-D2-Proto loop and the recurrent Proto loop in a reduced model comprising these two loops only. We vary the synaptic weights of the two loops *G*_*Proto−Proto*_ and *G*_*Proto−D*2_, run the simulation, and measure the frequency of stable oscillations in the firing activity (Fig. 2B). For low synaptic weights in either loop, the network does not leave the steady state. However, with increasing the *G*, the network is able to transition to the oscillatory regime (colored points in the phase space in Fig. 2B). Moreover, increasing the connection weight of the Proto-loop shifts the frequency of the network from the lower bound of *β* to the higher end of this band (Fig. 2C). Altogether, our results show that the experimentally-observed *β-*band frequency oscillations in the BG circuit could be resultant of the interactions between the pallidostriatal loops and a high-frequency oscillating di-synaptic loop.

### Oscillations frequencies in a realistic network model of BG circuits with experimentally-constrained time constants

The results of our rate model of BG circuits highlight the possible frequencies of oscillations driven by the various negative feedback loops of the network across a wide range of synaptic timescales and their possible interaction. However, neural dynamics in the rate model rely on a single time scale, ignoring the diverse timescales of neuronal and synaptic integration. The frequency of oscillations must thus be confirmed in a spiking neuronal network (SNN) where the membrane potential of the neurons is simulated using the leaky integrate-and-fire (LIF) neurons (see Materials and Methods for details). In this more detailed model, all timescales -the synaptic rise and decay time constants, neuronal integration time constant, and axonal delays-are extracted from experimental observations (see Table. 3-5 for values and references to the corresponding experimental studies). Moreover, the SNN model will allow us to both produce realistic activity patterns and to directly compare the phase-differences between the firing activity in the various neuronal populations during pathological *β* or artificially-induced oscillatory activity (as in (De la Crompe et al., 2020)). The investigation of the normal and pathological dynamics of BG activity in the SNN will be guided by the general results obtained by analytical derivation and simulations in the reduced rate model. An example of the spiking activity in a single neuron from each population is demonstrated in Fig. 3A (for presentation purposes, the APth is artificially set to 20 mV). After setting all the time constants and delays, we further verify the firing activity patterns of the network by mimicking the downstream effects of a short-term stimulation at the level of the motor cortex. Indeed, the temporal pattern of responses in the various neuronal populations of the BG is strongly coupled to the timescale of the synaptic and neuronal integration in the network. The effects of the cortical stimulation in all populations of our network thus offer an integrated view of the network’s temporal properties that can be directly compared to experimentally observed responses (Kita and Kita, 2011). These effects include a direct excitation in both the STN neurons as well as the D2-MSNs (Fig.3B). Both of these inputs contribute to shaping the GPe activity. Kita and Kita (2011) describe the GPe response as, primarily, an early excitation (from STN), then, an inhibition (from D2-MSNs), and lastly, a late excitation (again from STN) followed by a late long-inhibition. Since Proto neurons make up the majority of the GPe population (Mallet et al., 2012), we attribute this prototypical response to the Proto population. Our model reproduces similar tri-phasic activity in the Proto population activity. In particular, the relative timing of the onsets and peak in activity in the various populations of the model agree with the observed pattern. There are, however, some slight differences. Firstly, we do not observe the late and long-lasting inhibition in Proto observed by (Kita and Kita, 2011). This slow inhibition was shown to be driven mainly by GABA-b currents in Proto neurons, which are not included in our model. Since the time scale of GABA-b transmission is too slow (100-1000 ms decay (Gerstner et al., 2014)) to contribute significantly to the pathological *β-*band oscillations, we deliberately ignored this minor part of the inhibitory inputs to Proto. Secondly, the time scale of re-stabilization of Proto activity after the last excitation is somewhat faster than what is observed experimentally. The slow recovery of Proto neurons may be explained by their specific intrinsic electrophysiological properties that may not be captured in the LIF model. These differences are unlikely to play a significant role in the generation or modulation of pathological *β* oscillations and are thus beyond the scope of this study.

**Figure 3.**
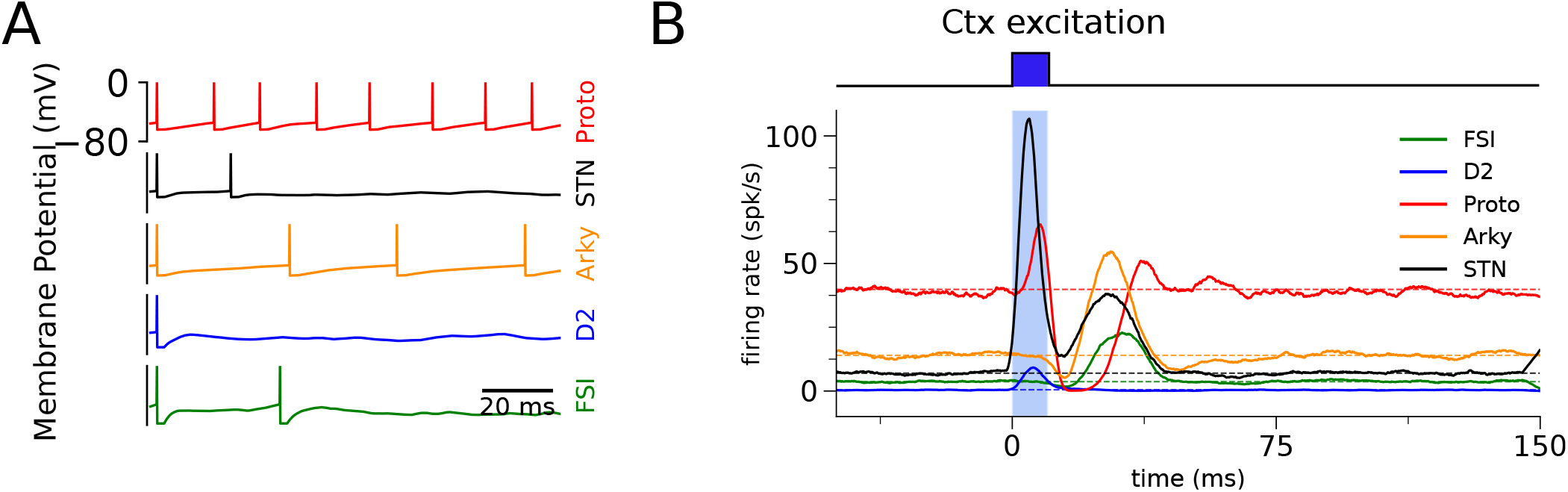
The spiking neural network model captures electrophysiological characteristics of the BG network. A) Example membrane potential traces of the LIF neurons in the model. The action potential height (APh) is artificially set to 0 mV for all neurons for presentation purposes. B) The effect of 10 ms cortex stimulation on the population activity of the neurons in the model. Each time course is an average of 10 different stimulation protocols.

Once all time constants and delays are set, we look at the frequency of the individual loops in the detailed SNN. We simulate each individual loop circuit when it is isolated from the network (Fig. 4A). In these simulations, the synaptic weights are set in a way to avoid the collective silencing of neurons in the population, while being sufficiently large to bring the circuit into the oscillatory regime (Fig. 4B). The temporal fluctuations in the mean activity across the neuronal populations reflect a clear oscillatory pattern while the high level of noise in the synaptic input induces large fluctuations within and between cycles, with waxing-and-waning oscillations in the mean activity. Importantly, the oscillations in mean activity are not reflected in regular spike trains firing at the oscillation frequency in single neurons (Fig. 4C). Rather, most neurons in the network see their probability of spiking oscillating in time but the synaptic noise and network heterogeneities make it difficult (if not impossible) to detect oscillations at the level of individual neuron spike trains. Due to the low firing rate of D2 neurons, their contribution to oscillations is even more difficult to tease apart at the level of spike trains, even when the firing activities of many neurons are displayed simultaneously (Fig. 4C). Nevertheless, oscillatory activity can be clearly revealed by spectral analysis of the population activity. To quantify the resultant frequencies, we run multiple sets of simulations (with varying parameter initialization) for each network and then average their corresponding power spectrum densities (Fig. 4D). The low-frequency component of the average population activity in our model may be reminiscent of local field potential recorded experimentally. The frequencies of oscillations generated in the SNN further confirm those of the rate model study. In particular, the recurrent connections between Proto neurons and the STN-GPe circuit generate *γ*-band frequencies (43 Hz for the STN-GPe circuit, 65 Hz for the Proto recurrent network). Whereas for realistic time constants, the two pallidostriatal loops oscillate with frequencies on the lower boundary of the *β-*band (Fig. 4D).

**Figure 4.**
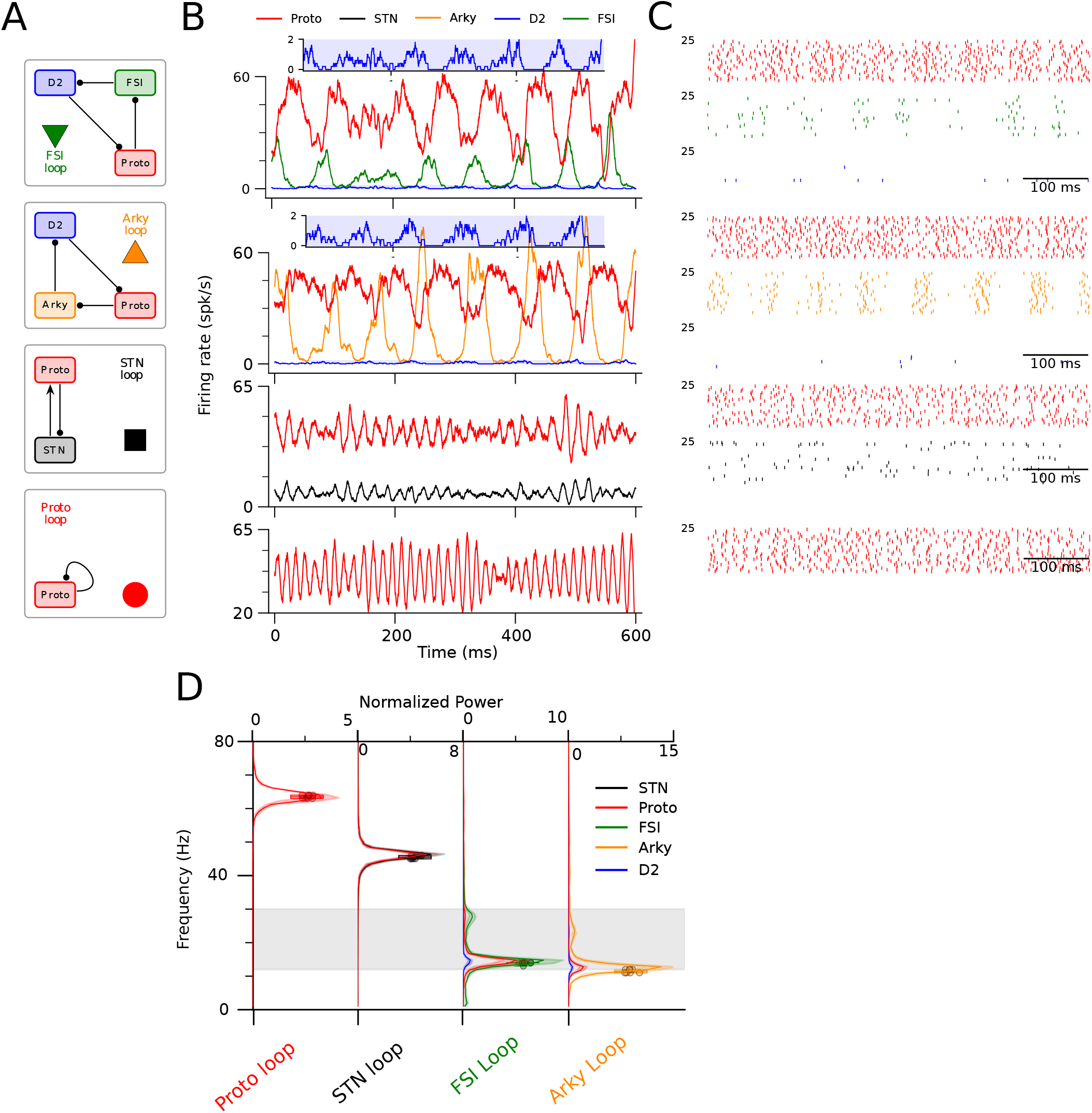
None of the individual feedback loops generate experimentally-observed *β-*band frequency oscillations. A) Schematics of individual feedback loops. B) Simulations of individual feedback loops yield their intrinsic frequency. Example average firing rate of *N* = 1000 neurons in each population of each given loop in (A). Insets highlights the activity of D2-MSNs where they exhibit oscillatory activity despite their low firing rate. C) Individual spikes of *N* = 25 randomly chosen neurons in (B). D) Normalized power spectrum density of the mean population firing rate in each of the loops in (A) for 8, 10-second long simulations (see Materials and Methods). The box plots summarize the peak frequencies (represented by circles) measured for the Proto population. The shaded area indicates the *β-*band (12-30Hz).

### The asynchronous healthy state

In the previous section, we artificially segregated the various feedback loop circuits from the BG network and increased the strength of the synapses to bring each sub-circuit into an oscillatory regime. However, for each subnetwork, mild synaptic weight does not lead to the emergence of oscillatory activity. For the complete BG network, these mild synaptic connections lead to an asynchronous state where all populations display non-oscillatory fluctuations driven by the fluctuations in their synaptic inputs (synaptic noise) and the relative timing of spike occurrence across the neuronal populations (Fig. 5A-B). This non-oscillatory state can be mapped onto the healthy state of the BG network, with the mean and fluctuations of external inputs adjusted to drive the observed distributions of firing rate in the various BG populations of anesthetized rats (De la Crompe et al., 2020; Mallet et al., 2005; Sharott et al., 2012; Sharott et al., 2017; Xiao et al., 2020). As the fluctuations in the network activity are non-oscillatory in this state, the cross-correlations between the spike trains of pairs of neurons of a given population are flat in all populations of the network (Fig. 5C, see also Fig. S5-1) and there is very low coherence between spike trains in a given population at any frequency (Fig. 5D, see also Fig. S5-1), as has been experimentally revealed in the BG neuronal population of animals in healthy condition (Mallet et al., 2008a). While most parameters of the model are constrained by experimental data, the synaptic weights are mostly free parameters of the model until this point. In the first part of the Results section, we have shown that mid-*β* oscillations can be driven by the interaction between several oscillating circuits in the rate model (Fig. 2B-C). Multiple loop circuits must thus be at play to produce the experimentally-observed *β* oscillations. Since the frequency of the resultant oscillations can change based on the contribution of each loop, it is now important to further constrain the synaptic weights in the model. While we cannot constrain all the synaptic weights with anatomical or physiological recordings, the weights along several connections of the circuits in the normal state have been set to reproduce the effect of artificial oscillatory drive by patterned optogenetic stimulation of the STN and Proto neurons in healthy rats reported in (De la Crompe et al., 2020), as explained thereafter.

**Figure 5.**
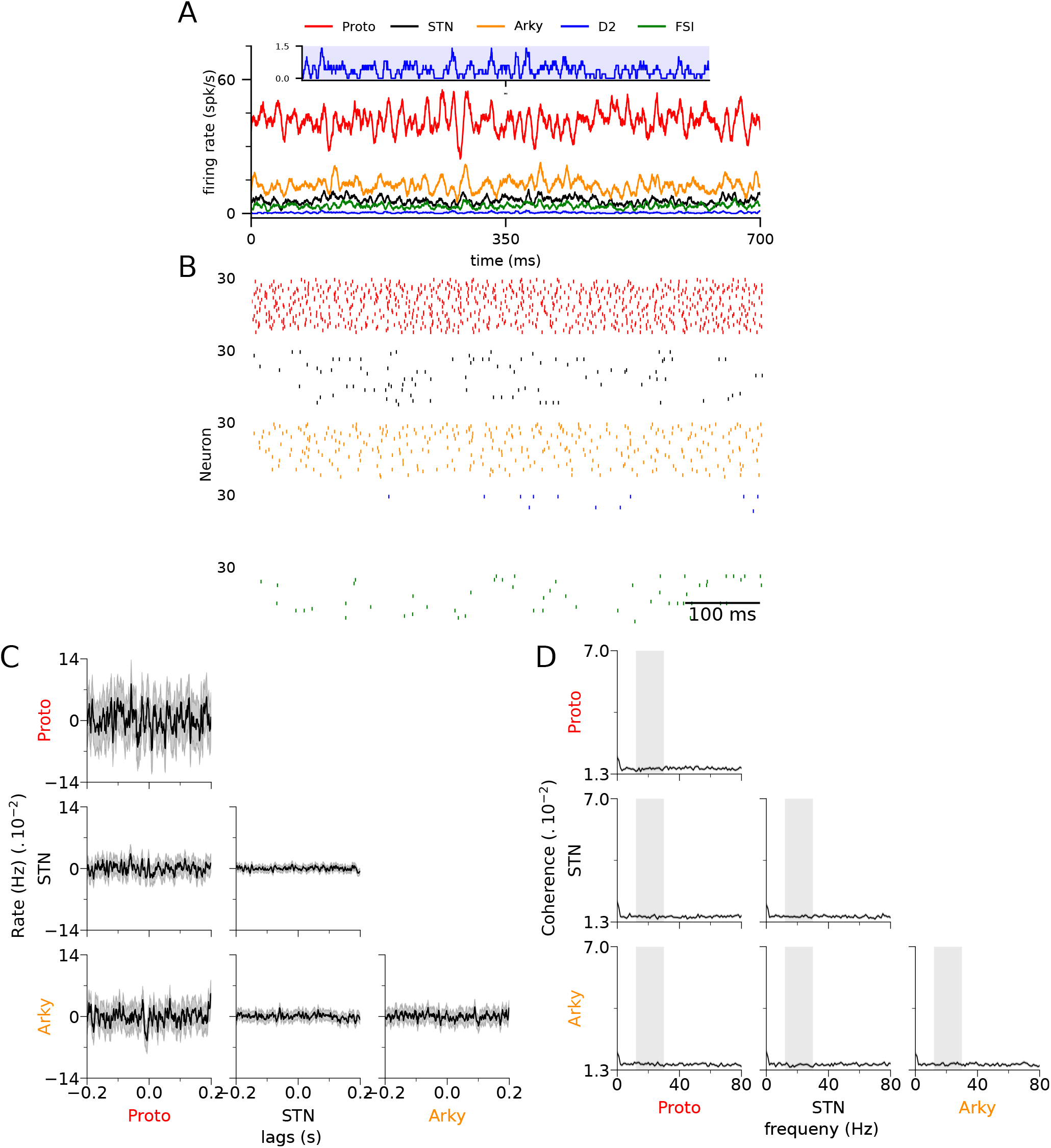
Healthy state of the tuned network does not exhibit any oscillatory activity. A) Average population firing rate time course of neurons simulated in an anesthetized healthy state. *N* = 1000 for all populations. Insets show the activity of D2-MSNs. B) Individual spikes of *N* = 30 randomly chosen neurons in (A). C) Average spike cross-correlation of 2000 randomly chosen pairs of neurons from the two populations specified by the rows and columns. No significant cross-correlation is observed on average among the pairs of populations. D) The average spike coherence of 1000 randomly chosen pairs of neurons from the two populations specified by the rows and columns. The network simulation is run for 25 seconds and the coherence is measured using 1-second windows. The shaded grey area indicates the *β* range (12-30Hz). No significant average spike coherence is observed on average among the pairs of populations.

### Activity modulation and phase differences during *β*-patterning

When *β* oscillations are artificially produced in healthy rats through the rhythmic activation of STN neurons with a 20 Hz half-sinusoidal optogenetic stimulation (Fig. 6A-left, (De la Crompe et al., 2020)), Proto neurons display a fast excitatory response following the STN activation with a short delay. The amplitude of the modulations of STN and Proto activity in these experiments reflects the strength of the direct excitation from STN neurons to Proto. Moreover, during the same protocol, Arky neurons are inhibited by the Proto, with a modulation whose amplitude reflects the strength of the inhibition from Proto to Arky. We set the synaptic weights between STN, Proto, and Arky neurons to reproduce the respective modulation of their activity during this experiment by delivering a 20 Hz half-sinusoidal depolarizing input to the STN neurons (Fig. 6A-right). Note that the relative phase difference between the various populations during this patterned stimulation is also accurately reproduced due to the appropriate time constants of the model derived from other experimental data (see Materials and Methods). In another experiment, the *β-*patterning of Proto neurons activity was induced in healthy rats through the non-specific rhythmic inhibition of GPe neurons with a 20 Hz half-sinusoidal optogenetic input that inhibits both the Proto and Arky populations (Fig. 6B-left, (De la Crompe et al., 2020)). We reproduce this patterned stimulation in the model with a rhythmic inhibitory input onto Proto adjusted to yield the same level of minimum activity in Proto neurons as in the experimental data (Fig. 3B-right). The relative amplitude of Proto and STN modulation in this experiment constrains the STN-Proto synaptic weight. Altogether, we are thus able to complete the tuning of the synaptic weights within the STN-GPe circuit and between GPe neuronal populations by replicating the *β-*patterning experiments. Note that as in Fig. 3B, the stabilization of Proto activity following a transient excitatory input from STN neurons is slower in the experimental data compared to the model. Again, this is likely due to the intrinsic cellular properties of Proto neurons that are not adequately captured by our LIF model. Nevertheless, the Proto activity in the model is stabilized by the end of each stimulation cycle resulting in evoked responses in STN neurons fully compatible with the experimental data both in terms of phase relationship and amplitude of the response (Fig. 6B). Note that pallidostriatal and Proto-Proto weights are also tuned in this experiment however this tuning will be later explained in the next sections.

**Figure 6.**
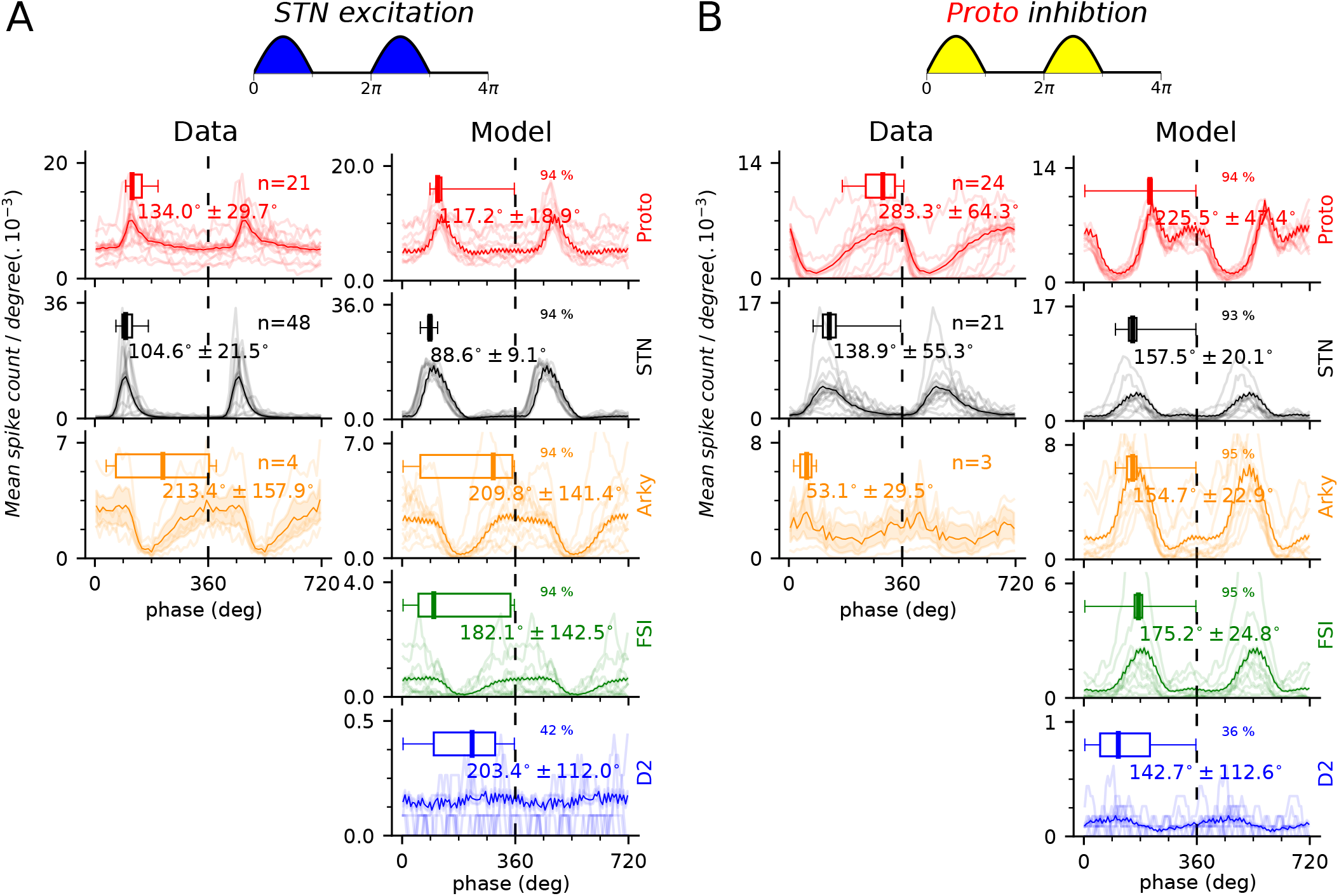
*β-*oscillation induction compared in the model and data. A) *β* oscillatory activity is artificially induced by exciting STN neurons at 20 Hz frequency as indicated in the schematic on top. Left: Single neuron spike linear phase histogram of Proto (red), STN (black), and Arky (orange) neurons during artificial *β-*patterning using non-specific opto-inhibition of GPe neurons in healthy rats using eArchT3 (re-adapted from De la Crompe et al. (2020)). Right: Single neuron spike linear phase histogram Similar to (left), but for our tuned model including the FSI (green) and D2-MSNs (blue) while applying similar excitatory input as the optogenetic manipulation on the left. B) Same as in (A) except for the fact that the *β* oscillatory activity is artificially induced by inhibiting Proto neurons at 20 Hz frequency as indicated in the schematic on top. In all panels, thinner lines represent randomly chosen individual neuron traces whereas solid line represents the average population phase histogram. The shaded area indicates the SEM. Box-and-whisker plots indicate median, first and third quartile, min, and max values. In the model, only the neurons (out of the total of *N* = 1000) that passed the Rayleigh test are taken into account, the percentage of which is indicated on the upper right quadrant of each panel.

### Dopamine–depletion (DD) in the model gives rise to *β* oscillations

The depletion of dopamine leads to pathophysiological and anatomical changes in the BG (Mallet et al., 2019). These changes are reflected both in changes in firing rate distributions (see Table. 6 for firing rates in the DD state) as well as some structural changes in the number and strength of synaptic connections in various BG pathways that are reported to occur after chronic DD. More precisely, (i) the number of connections in the projection from FSI to D2 neurons doubles (Gittis et al., 2011), (ii) the synaptic weights from Proto to STN neurons are doubled (Fan et al., 2012), and (iii) the synaptic weights of the recurrent inhibition between Proto neurons are doubled (Miguelez et al., 2012). We investigate the effect of DD in the BG by implementing all these functional and structural modifications in our BG network model. Following DD, the BG network sees the strength of the connections along several negative feedback loops strongly increased (doubled), namely the FSI-D2-Proto loop, the Proto-Proto loop, and the STN-Proto loop. The increased negative feedback along these loops is enough to push the network into an oscillatory regime. In this state, all the nuclei exhibit clear oscillatory activity that is observable in both the mean population firing rates as well as individual neurons’ spiking patterns (Fig. 7A-B). To characterize the oscillatory activity, we run multiple 25-second-long simulations of the network initialized at the DD state. We then use each simulation to derive the power spectrum densities (PSD) of the mean population firing rates (for more detail refer to Materials and Methods) and then average the PDSs over all repetitions. As shown in Fig. 7C, the average PSD exhibits a significant increase in the *β-*band oscillatory activity in all population after DD compared to the healthy state. The peak frequency of this oscillatory activity is 18 Hz. In addition, a small increase in the *γ*-band power (50-55 Hz) is observed in the Proto firing activity. The *γ*-band oscillatory activity in the Proto may give rise to the *β-γ* coupling observed in PD patients and in rats (López-Azcárate et al., 2010; Dejean et al., 2011). We confirm the relative role of each sub-circuit in the generation of the observed oscillatory activity by simulating each sub-circuit individually. With the synaptic weights of the DD state, and while disconnected from the rest of the network, the FSI-D2-Proto loop displays strong oscillatory activity in the low-*β* frequency, while the Proto loop and the STN-Proto loop are both close to the oscillatory regime with low oscillatory content in the gamma range (Fig. S7-2). Therefore, the pallidostriatal loop is the main driver of the observed pathological oscillations, and its interaction with either Proto-Proto or STN-Proto loop brings the oscillation in the mid-*β* range.

**Table 6.** Population firing rates (Hz) during anesthesia.

| Neuron | Anesthesia | Reference | Anesthesia DD | Reference |
| --- | --- | --- | --- | --- |
| D2-MSN | $0.49 \pm 0.14$ | (Sharott et al., 2017)<br>(Sharott et al., 2012) | $2.80 \pm 0.47$ | (Sharott et al., 2017) |
| STN | $7.0 \pm 5.8$ | (De la Crompe et al., 2020) | $24.5 \pm 11.8$ | (De la Crompe et al., 2020) |
| Arky | $14.1 \pm 8.3$ | (De la Crompe et al., 2020) | $12.2 \pm 6.0$ | (De la Crompe et al., 2020) |
| Proto | $39.8 \pm 13.7$ | (De la Crompe et al., 2020) | $21.6 \pm 12.2$ | (De la Crompe et al., 2020) |
| FSI | $3.67 \pm 3.22$ | (Mallet et al., 2005)<br>(Sharott et al., 2012) | 4.93 | (Xiao et al., 2020) |

Values are reported as mean ± SD. DD: Dopamine-depleted state.

**Figure 7.**
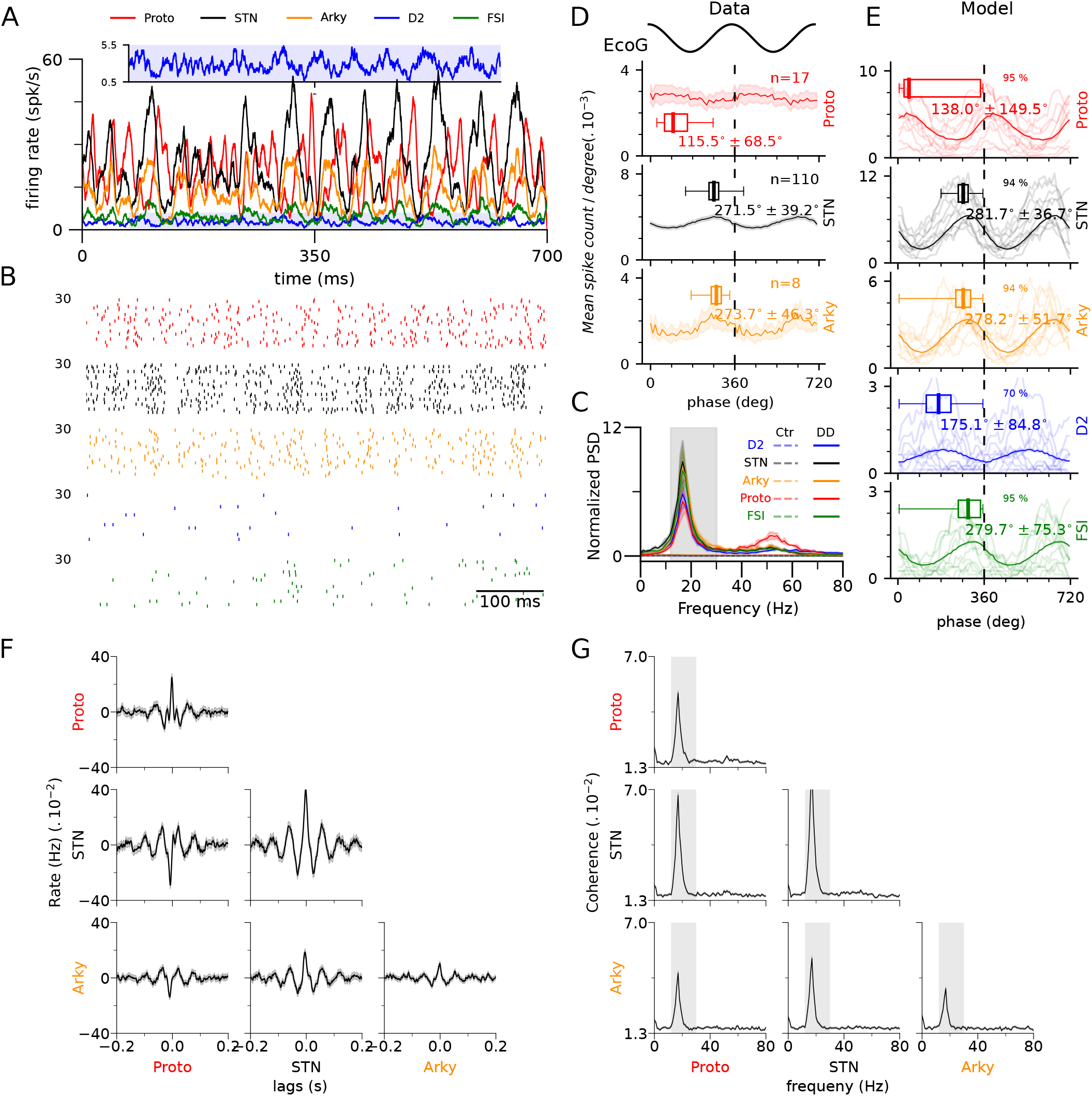
The model replicates the pathological phase locking and *β-*band oscillatory activity after transition to dopamine depletion. A) Average population firing rate time course of Proto (red), STN (black), Arky (orange), D2 (blue), and FSI (green) simulated in an anesthetized dopamine-depleted state. *N* = 1000 for all populations. Insets highlight the activity of D2-MSNs where they exhibit oscillatory activity despite their low firing rate. C) Normalized power spectrum of population average firing rate in (A). A significant peak is indicated in the *β-*band (12-30 Hz, shaded in grey). D) Single neuron spike linear phase histogram of Proto (red), STN (black), and Arky (orange) neurons during spontaneous *β* oscillations aligned to the recorded EcoG in anesthetized parkinsonian rats (re-adapted from De la Crompe et al. (2020)). E) Single neuron spike linear phase histogram Similar to (D), but for our tuned model including the FSI (green) and D2-MSN (blue). Phases are aligned to Proto peaks after *β* filtering. Thinner lines represent results for randomly chosen individual neurons. The solid line represents the average population phase histogram and the shaded area indicates the SEM. Box-and-whisker plots indicate median, first and third quartile, min, and max values. In the model, only the neurons (out of the total of *N* = 1000) that passed the Rayleigh test are shown, the percentage of which is indicated on the upper right quadrant of each panel. C) Average spike cross-correlation of 2000 randomly chosen pairs of neurons from the two populations specified by the rows and columns. All population pairs exhibit significant spike cross-correlation with lags corresponding to the *β-*band frequencies. D) The average spike coherence of 1000 randomly chosen pairs of neurons from the two populations specified by the rows and columns. The network simulation is run for 25 seconds and the coherence is measured using 1-second windows. The shaded grey area indicates the *β* range (12-30Hz). All population pairs exhibit enhanced spike coherence in the *β* range.

While the frequency of the oscillatory activity generated in our BG network in the DD state agrees with abnormal *β* - band oscillations, the nature of the interactions between the different populations that give rise to the *β* oscillations can be revealed by comparing relative phase distributions of spikes across all populations of the network. Indeed, De la Crompe et al. (2020) have reported the individual spike phase histogram of the Proto, STN, and Arky neurons relative to the motor cortex electrocorticogram (ECoG) signal in parkinsonian rats (re-adapted in Fig. 7D). Their results provide a great biological reference for us to validate our model with. To compute the relative spike phase histograms, we use the Proto mean population firing rate as our reference instead of the ECoG signal, as We do not include the cortex in our model. Using 25-second long simulations, we then calculate the relative phases of all the spikes for each individual neuron separately. In order to be able to compare the phase distribution to those of the experimental data, we make use of the findings of Sharott et al. (2017) where they show that in parkinsonian rats, the D2-MSNs exhibit anti-phase activity relative to the recorded ECoG. Therefore, we shift the phase distributions of all neurons so that the average phase distribution of D2-MSNs is anti-phase to the *β-*filtered ECoG signal in Fig. 7D. The final results of the phase distributions for all the populations in the DD state are shown in Fig. 7E. Our results show that the phase relationships between the Proto, STN, and Arky neurons in the model closely resemble those of the experimental data (Fig. 7D-E). In particular, peak STN activity is in phase with Proto inhibition, consistent with a main oscillatory drive coming from the GPe rather than from STN (see Fig. 6). The phase difference between Proto and STN argues against a significant contribution of the STN-Proto loop to the DD state oscillations observed here. To confirm the marginal (if any) role played by the STN-Proto loop, we also replicate the effect of manipulations of GPe and STN activity performed experimentally (see below). To our knowledge, the phase relationship of FSIs in the DD state has not yet been reported experimentally. As the FSI-D2-Proto sub-circuit plays a central role in the generation of the observed oscillatory activity, our model makes a strong prediction concerning the relative phase distribution of the spiking activity of FSI neurons relative to other BG populations that may be tested in future experiments.

In the DD state, the strong *β-*band oscillatory activity of the network can be revealed by cross-correlation analysis of the spiking activity of the Proto, STN, and Arky neurons (Fig. 7F). Indeed, the cross-correlation analysis shows a strong oscillatory pattern of correlation between neurons in a given population, as well as strong synchrony between the different populations. The peak and trough of the cross-correlations at zero-lag also indicate the phase or anti-phase entrainment between the populations, confirming the previously presented phase-difference analysis, with Arky and STN neurons oscillating in phase and Proto neurons displaying an anti-phase relation to each of them. This *β-*entrained correlation is evident in all the pairs of populations except for among the D2 neurons, due to their very low firing activity levels (Fig. S7-1). The oscillatory synchronization between neurons within and across BG neuronal populations can also be revealed by the strongly increased coherence in the *β-*band between spike trains (Fig. 7G, Fig. S7-1).

### Proto inhibition, and not STN, abolishes *β*-band oscillations in dopamine-depleted state

STN-GPe network has been a long-standing candidate for explaining the origins of the pathological *β* oscillations in the BG (Plenz and Kital, 1999; Terman et al., 2002; Nevado Holgado et al., 2010; Kumar et al., 2011; Pasillas-Lépine, 2013; Nevado-Holgado et al., 2014; Pavlides et al., 2015; Shouno et al., 2017; Koelman and Lowery, 2019; Chen et al., 2020). However, De la Crompe et al. (2020) have recently shown that with collective opto-inhibition of the STN neurons, the *β* oscillations are still maintained at the level of both the cortex and GPe. In light of their results, it could be debated that the STN-loop is not the generator of the *β* oscillations after all. In order to test this hypothesis, we inhibit the STN neurons using a uniform hyperpolarizing input that is applied to all the neurons in STN (Fig. 8A). This rather large inhibition does not lead to any significant change in the average firing rate of Proto neurons. This lack of modulation indicates that the synaptic feedback from STN to the Proto population is rather weak. We inhibit the network for 25-seconds and use the average population activity time courses of each population to measure their respective PSDs. Surprisingly, the STN inhibition not only does not diminish *β* oscillations in other populations, but it also boosts the oscillation power in the *β-*band (Fig. 8B). On the other hand, understandably, the reduction in STN activity due to the inhibition results in a decrease in the power of the *β-*band oscillations (Fig. 8B).

**Figure 8.**
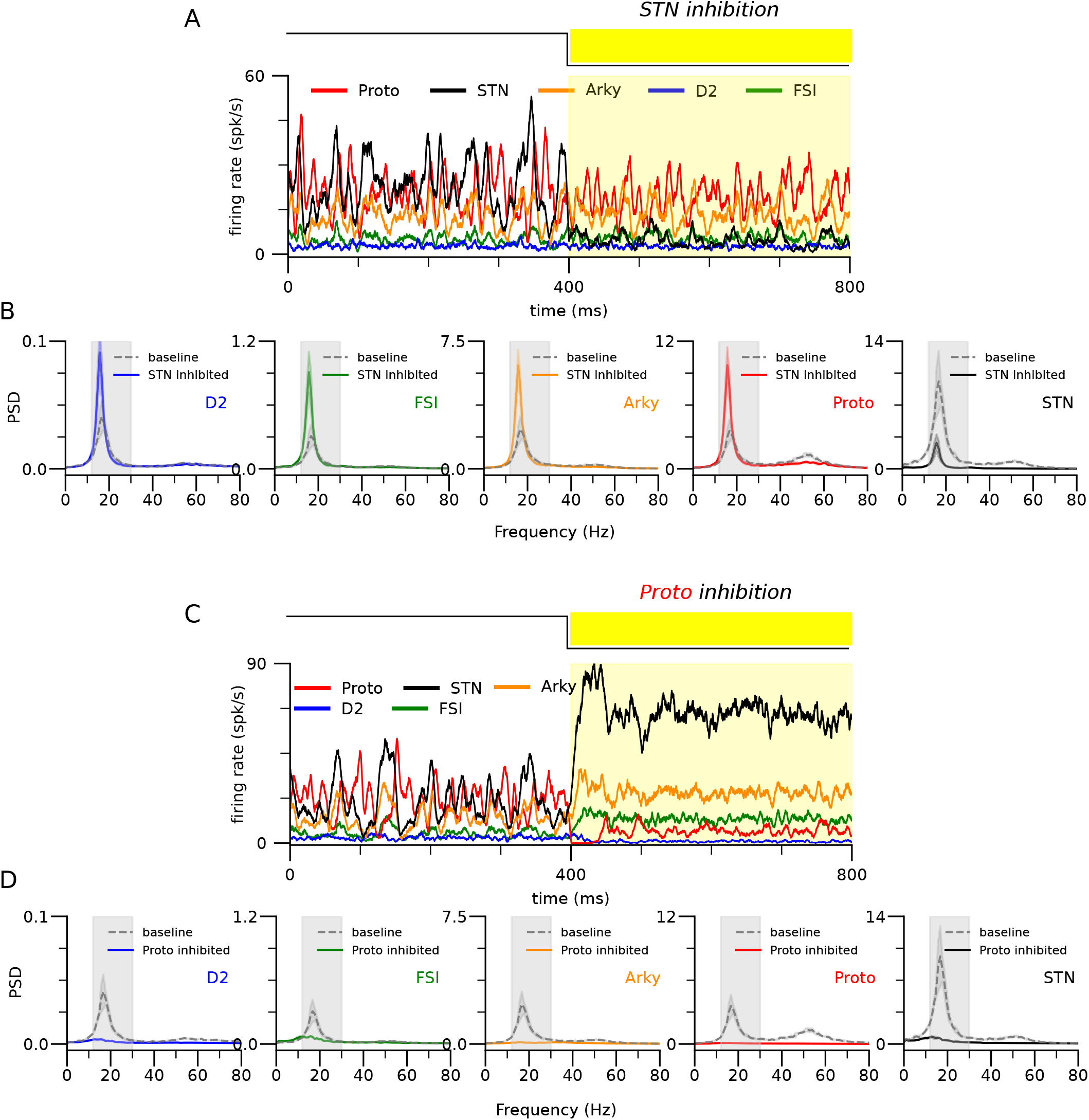
Proto inhibition, and not STN, abolishes *β-*band oscillations in dopamine depleted state. A) Average population firing rates in a network of Proto, STN, Arky, D2, and FSI when in a dopamine-depleted state (DD) the STN neurons are inhibited. The yellow shaded color indicates where external inhibitory input is applied to STN. B) Power spectrum densities (PSD) of each of the populations compared between baseline in DD (solid colored) and when inhibition is applied to STN (dashes grey). STN inhibition does not uplift *β-*oscillation. C) Average population firing rates in a network of Proto, STN, Arky, D2, and FSI when in a dopamine-depleted state (DD) the Proto neurons are inhibited. The yellow shaded color indicates where external inhibitory input is applied to STN. B) Power spectrum densities of each of the populations compared between baseline in DD (solid colored) and when inhibition is applied to Proto (dashes grey). Contrary to STN, Proto inhibition abolishes *β-*oscillation in all populations. Grey shaded area indicates the *β* range (12-30Hz).

In a similar experiment as above, De la Crompe et al. (2020) targeted the whole GPe population using a non-specific inhibitory opsin. The Opto-inhibition of the GPe neurons abolished the *β-*oscillation in activities of both the motor cortex ECoG and STN activity. Their results suggest a key central role for the GPe in the generation of the *β* oscillations. Therefore, we hypothesize that since the Proto population is the shared node in all our loops, its inhibition should in principle abolish all oscillatory activity in the DD state. We then tested this hypothesis by inhibiting all the Proto neurons using a uniform hyperpolarizing input. Contrary to STN, Proto inhibition significantly affects the STN activity via disinhibition. This is partly due to the increase in STN-Proto weight after dopamine depletion (Fan et al., 2012)(Fig. 8C). The power spectrum density analysis shows that the Proto inhibition results in total suppression of *β-*oscillation in the activity of all the populations as shown in Fig. 8D. These findings rule out the significant contribution of the STN-loop in the generation of *β* oscillations. In fact they support the hypothesis that the Proto–central loops are orchestrating the *β* oscillations in the DD state.

## Discussion

Building on up-to-date experimental evidence, our model shows that the neuronal and synaptic properties of dopamine-depleted pallidostriatal circuits can fully explain the *β* synchrony observed in experimental parkinsonism. The loops linking pallidal (Arky and Proto) and striatal (FSI and D2) neuronal populations generate oscillations in the low-*β* range (*<* 15 Hz). However, the interactions between these loops and strong inhibitory reciprocal connections within the pallidal Proto neurons can increase the oscillatory frequency in the complete network, driving abnormal *β* oscillations with experimentally-observed frequencies (≈ 20 Hz). Consistent with data, inhibition of the GPe, but not the STN, impairs the expression of abnormal *β* oscillations (De la Crompe et al., 2020). Indeed, the two GPe neuronal populations play a crucial role in the generation of pathological oscillatory activity in our model. These neurons participate in the two pallidostriatal reciprocal and recurrent circuits which induce *β* oscillations after dopamine depletion. The GPe having a central role in the circuitry also imposes phase relationships between various BG populations that are consistent with those reported in parkinsonian animals and when *β* oscillations are artificially driven through GPe activity modulation (De la Crompe et al., 2020). On the contrary, the timescale of activity propagation in the STN-GPe network is inconsistent with a putative generation of *β* rhythms and the observed strength of its reciprocal connections is too low to be the main drive of oscillatory activity (Fig. S7-2).

The large amount of data collected in parkinsonian rats, including the phase relationships between BG neuronal populations during *β* oscillations and the effects of neuronal manipulation on abnormal *β*, constrains the underlying mechanisms as shown here. We thus chose to build our model according to the data collected from normal and parkinsonian rats (and, when unavailable, from mice). Future experiments uncovering phase relationships between BG neuronal populations during abnormal *β* oscillations in patients or non-human primates (NHPs) could inform us about a possible parallel in the *β-*generation mechanisms between parkinsonian patients, NHPs, and rats. Parkinsonian rats display abnormal *β-*activity that is highly behavior and state-dependent (Mallet et al., 2008b; Avila et al., 2010; Brazhnik et al., 2021) and can spread from mid-*β* (15-25 Hz) under anesthesia (Mallet et al., 2008b; De la Crompe et al., 2020) to high-*β* (25-30 Hz) in awake behaving parkinsonian rats (Bergman et al., 1990; Avila et al., 2010; Brazhnik et al., 2014; Delaville et al., 2015; Brazhnik et al., 2016). The interaction between multiple oscillation generation circuits, as proposed here, naturally accounts for frequency changes as the relative strength of each oscillatory circuit can influence the frequency of global oscillations within the *β* range (Fig. 2C).

The mechanism for abnormal *β* oscillations revealed here may also be at play in parkinsonian patients and in other animal models. However, the properties of abnormal *β* activity differ between animal models and different generation mechanisms may be involved. Firstly, the frequency of abnormal *β-*oscillations under dopamine depletion varies across species (Stein and Bar-Gad, 2013; Smith and Galvan, 2018). Parkinsonian mice mostly show increased field potential activity in the delta range when active and are not reported to exhibit exaggerated *β* activity (Willard et al., 2019; Whalen et al., 2020) (but see (Baaske et al., 2020)). NHP models of PD display excessive *β* with peak frequencies in a reduced low-*β* range (Nini et al., 1995; Raz et al., 2001; Stein and Bar-Gad, 2013; Deffains and Bergman, 2019). In PD patients, the frequency of abnormal *β* activity varies across individuals (Kühn et al., 2009), with local field potential recordings typically showing exaggerated *β-*range activity with frequencies within the low (10-15Hz) and/or high (20-30Hz) bands (Hutchison et al., 2004), possibly reflecting different generation mechanisms. Secondly, previous experiments suggest that STN plays a central role in abnormal *β* generation in NHP (Deffains and Bergman, 2019; Tachibana et al., 2011) while it does not in parkinsonian rats (De la Crompe et al., 2020). A circuit involving the STN, such as STN-GPe loop or the hyperdirect loop (Leblois et al., 2006; Pavlides et al., 2015), may thus contribute to the low-*β* observed in NHP. A recent modeling study suggests that STN-GPe and pallidostriatal circuits could interact to drive such abnormal *β* oscillations (Ortone et al., 2022). The multiple bands of *β* activity observed in PD patients may reflect a combination of mechanisms only partially reflected in the various animal models. Alternatively, difference in the relative strength of the pallidostriatal and Proto loops leading to oscillation frequencies (see Fig 2C) may account for different frequencies between patients.

While pallidostriatal circuits can generate pathological *β* oscillations showing characteristics of those observed in parkinsonian rats, we cannot exclude other circuits taking part in *β* generation. In particular, the large BG-thalamo-cortical loop circuits (Leblois et al., 2006; Pavlides et al., 2015), or the STN-GPe loop may interact positively with BG-generated *β* oscillations (Corbit et al., 2016; Ortone et al., 2022; West et al., 2022). However, experiments in parkinsonian rats clearly show that the cortex and STN are not necessary for the expression of *β* oscillations while in fact GPe is (De la Crompe et al., 2020). It leaves a possibility for BG-thalamic loops (Str-GPe-SNr-Thal-Str) to interact with the presently investigated circuits (Brazhnik et al., 2016). GPe Arkypallidal neurons project to striatal FSIs, an anatomical connection not considered in our model. It could oppose the negative feedback in the Arky-MSN-Proto loop. However, This connection is considerably weaker than the projections of Proto onto FSI (Corbit et al., 2016) and is thus unlikely to impede or significantly alter oscillation generation. Within the pallidostriatal circuit, interactions between Proto and Arky may also participate in fast oscillations generated by the GPe alone (Gast et al., 2021).

Our model makes clear predictions concerning the phase of BG populations during *β* oscillations that can be tested both in rodents and in NHP. On one hand, the phase of FSI firing, yet unknown in rodents and a critical player in our model, may motivate future experiments in parkinsonian rats. On the other hand, the phase relationships between STN neurons and the two GPe populations recently revealed in NHPs (Katabi et al., 2022) can be quantified in future studies to test whether similar mechanisms are at play in this animal model. Synchronization at 10-13Hz emerges in the BG-thalamo-cortical network of GAERS rats during spike-and-wave discharges (SWD) characteristic of absence epilepsy seizures (Paz et al., 2005). The phase relationship between STN and GPe neurons in SWD is similar to their phase relationship during parkinsonian *β* oscillations (Paz et al., 2005; De la Crompe et al., 2020). The pallidostriatal loop, which can produce low-*β* rhythm as shown here, may also be involved in such pathological oscillatory activity, possibly amplified or complemented by the larger hyper-direct loop (Arakaki et al., 2016). Dopamine depletion induces a number of modifications in the BG network, some of which are likely causal to the emergence of abnormal oscillatory activity (Quiroga-Varela et al., 2013).

Our model incorporates functional (e.g. changes in excitability) and structural (remodeling of synaptic connectivity) changes reported in the pallidostriatal and pallidosubthalamic circuits. In particular, the number of synapses from FSI to D2 (Gittis et al., 2011) as well as the strengths of the synapses in STN-GPe and GPe-GPe connection are increased (Fan et al., 2012; Miguelez et al., 2012). Unlike immediate functional changes induced by the lack of dopamine (e.g. changes in D2 excitability, or the modulation of cortico-striatal synaptic transmission), these structural changes rather appear over days and are regarded as homeostatic processes that compensate for immediate activity changes induced by dopamine loss (Gittis et al., 2011; Ketzef et al., 2017; Villalba and Smith, 2018). Correspondingly, the observed hyperactivity of D2-MSNs after dopamine depletion and predicted by the classical model of PD (Albin R., 1989) are reversed over days following DD (Parker et al., 2018), possibly due to the increased inhibition from FSI after synaptic remodeling. The emergence of pathological oscillatory activity is induced in our model by the structural changes in the D2-FSI connections and is thus expected to arise with a similar delay following immediate dopamine depletion. This is consistent with the relatively late emergence of pathological *β* oscillations in these models (Leblois et al., 2007; Degos et al., 2009; Quiroga-Varela et al., 2013).

Our modeling study provides a candidate mechanism for the generation of *β* abnormal activity in PD, compatible with evidence in parkinsonian rats, and makes clear predictions to be tested both in rats and other animal PD models. The variety of properties of *β* oscillations observed in PD patients and animal models (frequency band, phase relationships) may reflect very different generation mechanisms. As the link between abnormal synchronizations of neuronal activity and abnormal movement in PD is still controversial (Kühn et al., 2006; Leblois et al., 2007; Quiroga-Varela et al., 2013; Sharott et al., 2014; Neumann et al., 2016), it is crucial to uncover the neural mechanisms for the generation of oscillatory activity in each condition and context to improve our understanding of the pathophysiological mechanisms underlying PD and develop new therapeutic strategies.

## Appendix. 1 Theoretical analysis

In this appendix, we aim to analytically derive the frequencies of the oscillatory instabilities for individual isolated loops of the model network. For the sake of simplicity, we consider an all-to-all connectivity matrix *J*^*βα*^ = *I*. In this case, the steady states of the network in response to constant inputs can be obtained by solving the fixed-point of Eq. 2. For steady states where all neuronal populations have inputs above the threshold, the activities are determined by a set of linear equations that can be solved straightforwardly. The conditions for the stability of steady states can then be derived. Indeed, a fixed-point solution of the dynamical equations is stable if any small perturbation around it eventually decays at large time. If certain perturbations increase with time, the fixed point is unstable. To investigate the stability of a steady state, we study the equations of the dynamics linearized around that state (Strogatz, 2018). We consider a perturbation term for the output *m*^*βα*^ described in Eq. 2.

**Figure S2-1.**
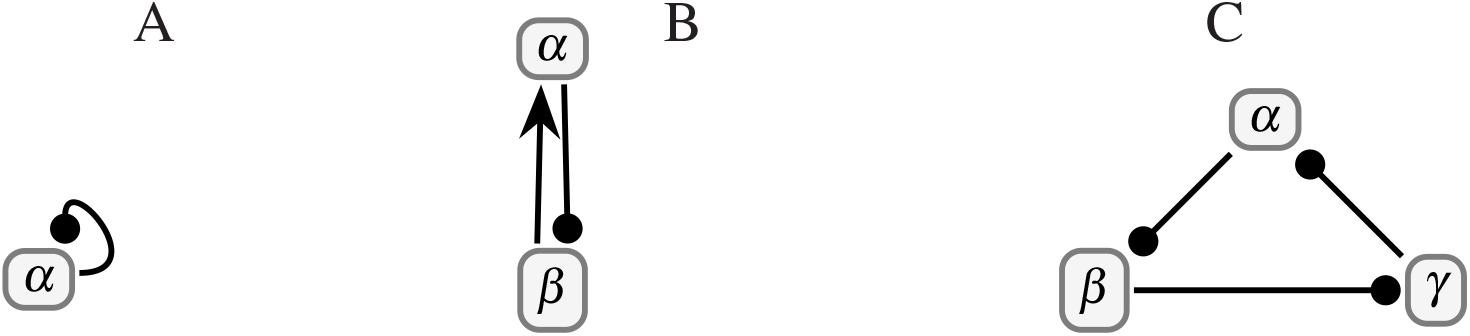
Theoretically-investigated possible oscillation-generator networks. A) A self-inhibiting population. B) A reciprocal recurrent circuit comprising an excitatory and an inhibitory population. C) Three populations creating a loop with inhibitory connections.

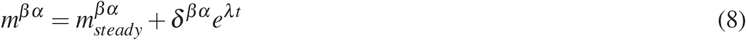

replacing this in Eq. 2 we get

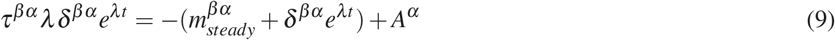

Then, for an activation function with unit gain, we have *A*^*α*^ = *I*^*α*^. Therefore if we consider a network of only two populations *α* and *β* (Fig. S2-1B), from Eq. 2 we have

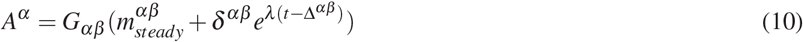

At steady state, from Eq. 2 we can write 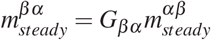. Therefore, replacing *A*^*α*^ in Eq. 9 we have

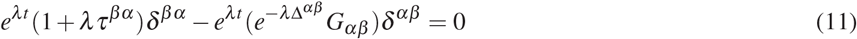

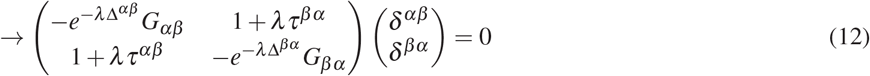

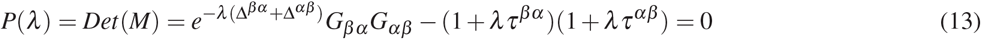

The solutions to this equation are in general complex numbers. The steady state is stable provided that *Re*(*λ*) *<* 0 for all the solutions. It is unstable if at least one solution with *Re*(*λ*) *>* 0 exists. If, for this solution, *Im*(*λ*) *<* 0, the system undergoes a non-oscillatory instability. If *Im*(*λ*) *>* 0, the instability is a Hopf bifurcation (Strogatz, 2018) at a frequency *Im*(*λ*). For such an oscillatory instability *λ* = *iν* we get:

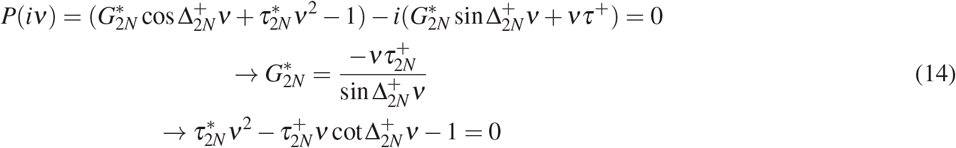

Where

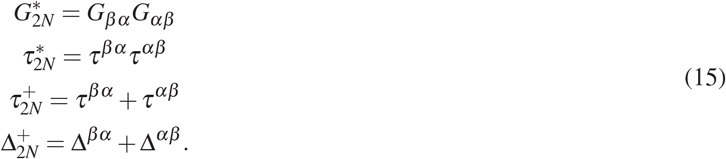

In particular,we can derive the resultant frequency for a feedback loop with three nodes (Fig. S2-1C) (i.e Proto-FSI/Arky-D2) as follows:

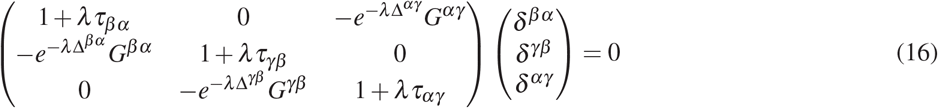

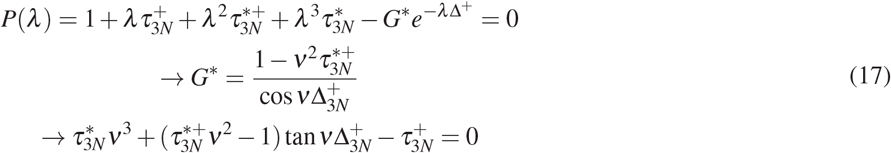

Where

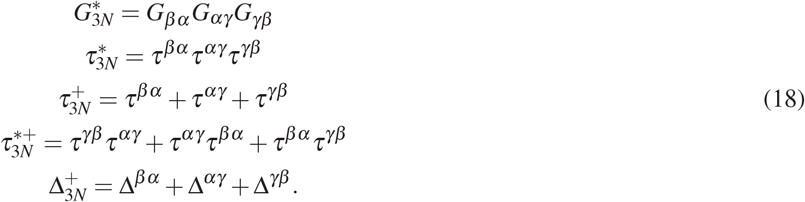

We can also calculate the frequency of oscillation for a neuron that contacts itself (Fig. S2-1A) to mimic the Proto lateral inhibition

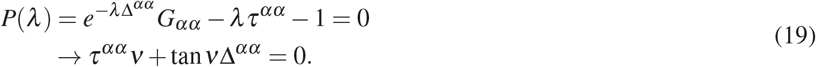

In order to estimate the oscillation frequencies of each of the individual loops we have solved the corresponding characteristic equations using *scipy*.*optimize*.*fsolve* function in python (Fig. 2A).

## Appendix. 2 Derivation of *K*_*βα*_

Here we calculate the number of convergent neurons from the presynaptic population *α* onto the postsynaptic population *β, K*_*βα*_. The values for the number of neurons in each population used in the calculations are taken from Table. S1-1. We assume an average of 3 boutons per synapse is a reasonable number.

- **D2-FSI**: As previously mentioned by (Guzmán et al., 2003); There are 2840 striatal neurons inside the volume of a spiny dendritic tree (Oorschot, 1998; Kincaid et al., 1998), 5% of which are FSIs *n* = 140 (Koós and Tepper, 1999). In addition, (Gittis et al., 2011) have discovered that the connection probability of recorded pairs of D2-MSNs and FSI are 36% within a radius of 250*µm* radius. Therefore we estimate that the biological number of FSI neurons converging to a single D2-MSN is given by

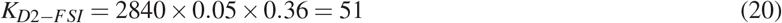
- **FSI-Proto**: In a study, Bevan (1998) have shown that 6/23 labeled pallidal neurons projected to the neostriatum where the average number of boutons of each is 791 (ranging from 329 to 1353). They have also shown that 19 to 66% (average from Table. 3 is 44 %) of these boutons make contact with the PV+ interneurons (Bevan, 1998). Assuming a minimum of 3 boutons converging onto postsynaptic neuron (Guzmán et al., 2003), yields,

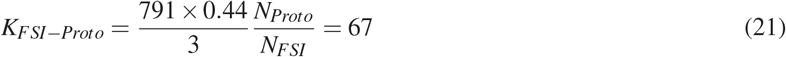
- **D2-Arky**: The projections from Arky to the striatum are rather understudies and experimental data on the number of projections initiating from Arky are scarce. Therefore, assuming that the projections from Arky to striatum are much more sparse than projections from Proto, we roughly estimate *K*_*D*2*−Arky*_ *≈* 10.
- **Proto-D2**: The number of boutons in the GPe formed by an axon of an indirect projection neuron is 226 (Koshimizu et al., 2013). Therefore,

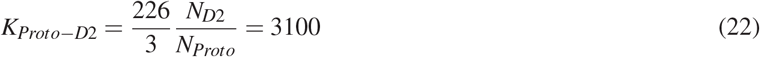
- **STN-Proto**: An individual GP neuron forms about 275 synapses in the STN (Baufreton et al., 2009). This would yield,

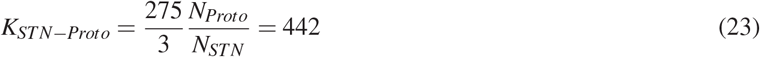
- **Proto-STN**: 8/10 STN neurons studied contributed projections to the GPe, and each of these made between 245 and 1450 presynaptic boutons (average of 558 for the eight cells with GPe projections) (Koshimizu et al., 2013). Therefore similar to STN-Proto maximal number of convergence is

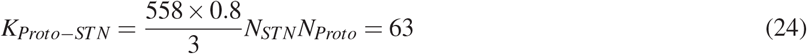
- **Proto/Arky-Proto**: The neurons in the border region had an average of 264 boutons, and those in more medial regions had an average of 581 boutons (Sadek et al., 2006). Which on average gives 422 boutons.

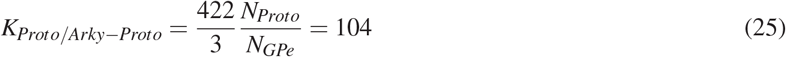

**Supplementary Table S1-1.**
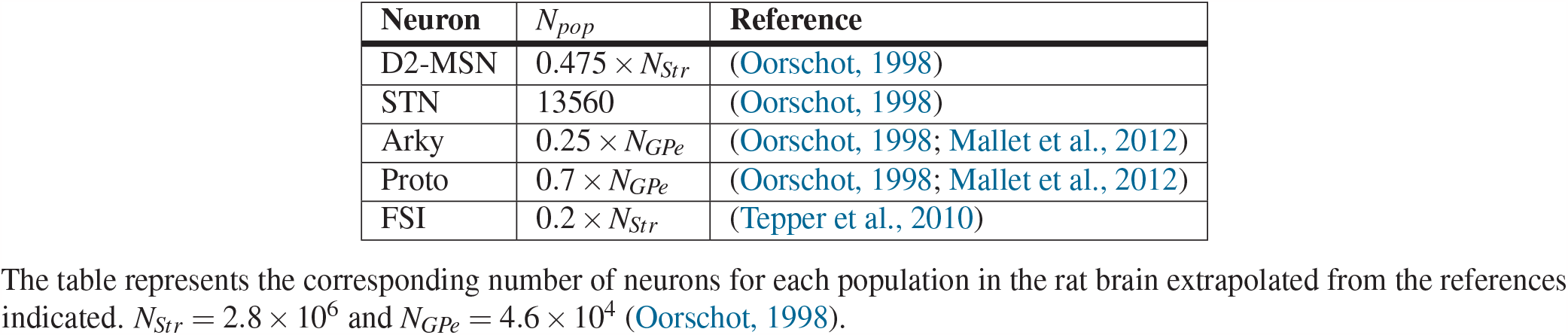
Number of neurons in the neural populations.

| Neuron | $N_{pop}$ | Reference |
| --- | --- | --- |
| D2-MSN | $0.475 \times N_{Str}$ | (Oorschot, 1998) |
| STN | 13560 | (Oorschot, 1998) |
| Arky | $0.25 \times N_{GPe}$ | (Oorschot, 1998; Mallet et al., 2012) |
| Proto | $0.7 \times N_{GPe}$ | (Oorschot, 1998; Mallet et al., 2012) |
| FSI | $0.2 \times N_{Str}$ | (Tepper et al., 2010) |

The table represents the corresponding number of neurons for each population in the rat brain extrapolated from the references indicated. $N_{Str} = 2.8 \times 10^6$ and $N_{GPe} = 4.6 \times 10^4$ (Oorschot, 1998).

**Supplementary Figure S5-1.**
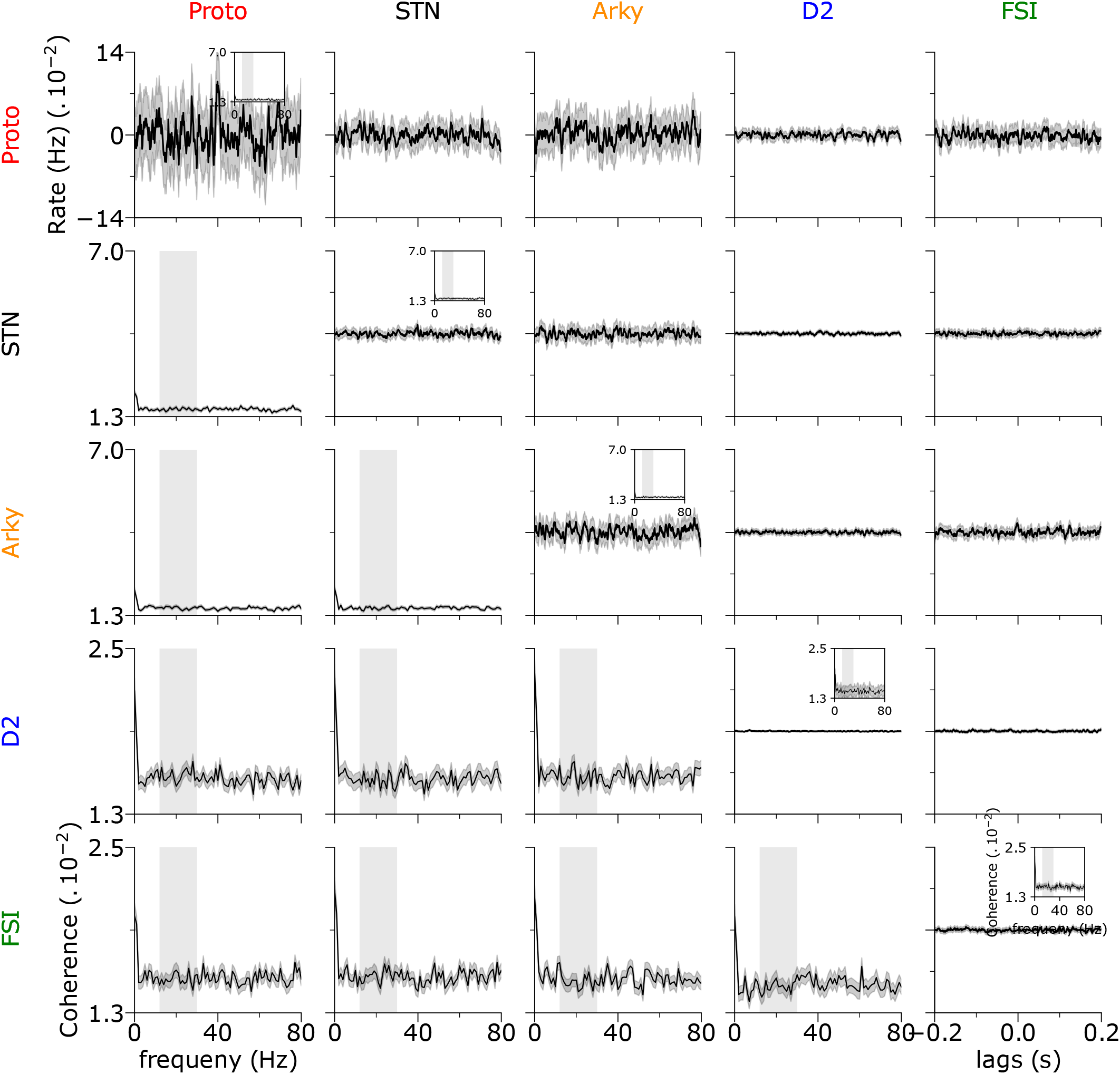
Cross-correlation and Coherence of spikes in the healthy state. Below the diagonal: The average spike coherence of 1000 randomly chosen pairs of neurons from the two populations in rows and columns. Each neuron is simulated for 25 seconds and the coherence is measured by using 1-second long windows. Almost all populations exhibit significant coherence in the *β-*band. The shaded grey area indicates the *β* range (12-30Hz). Diagonal and below: Average spike cross-correlation of 2000 pairs of neurons. Insets on the diagonal show spike coherence.

**Supplementary Figure S7-1.**
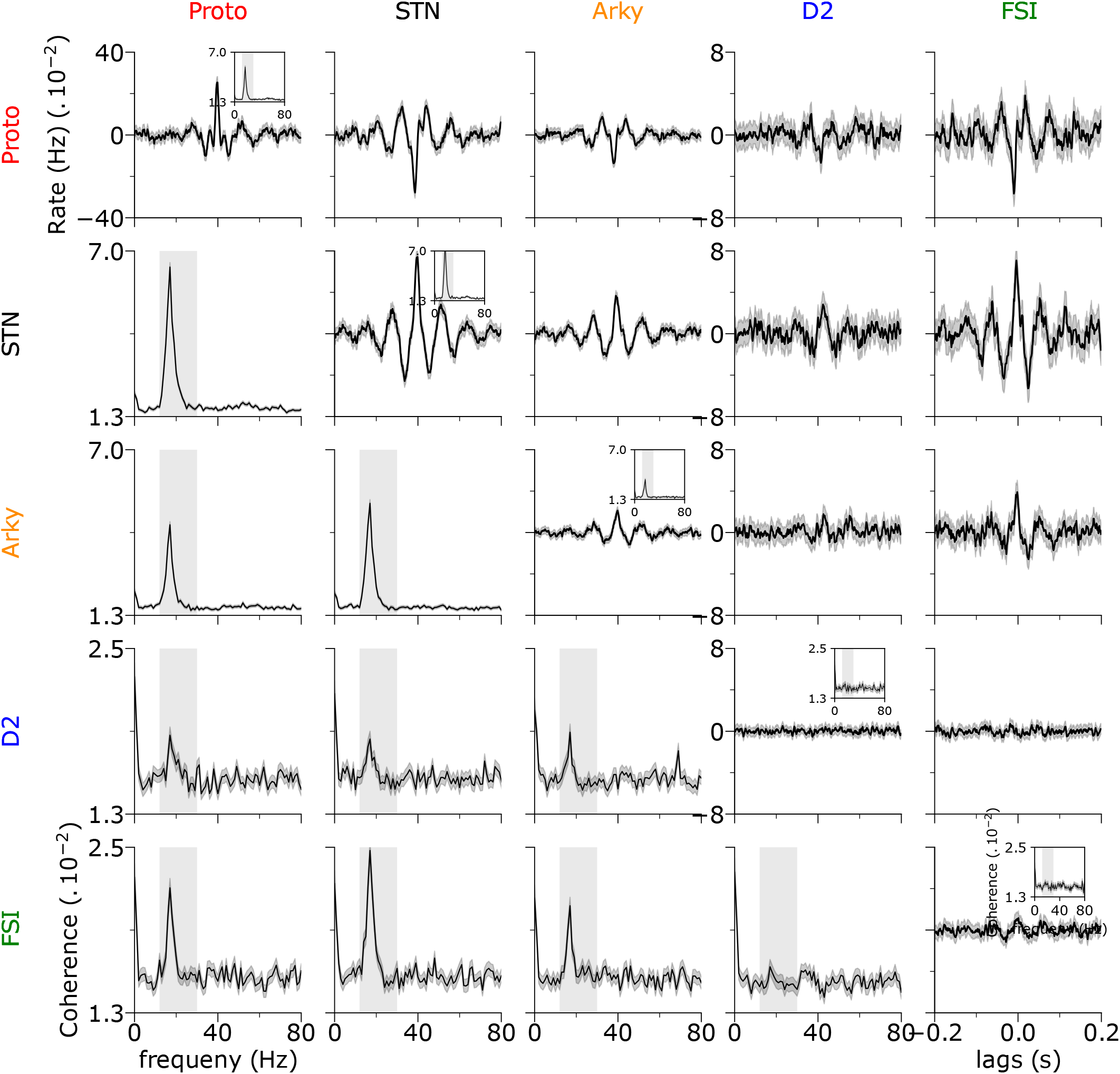
Cross-correlation and Coherence of spikes in dopamine depletion. Below the diagonal: The average spike coherence of 1000 randomly chosen pairs of neurons from the two populations in rows and columns. Each neuron is simulated for 25 seconds and the coherence is measured by using 1-second long windows. Almost all populations exhibit significant coherence in the *β-*band. The shaded grey area indicates the *β* range (12-30Hz). Diagonal and below: Average spike cross-correlation of 2000 pairs of neurons. Insets on the diagonal show spike coherence.

**Supplementary Figure S7-2.**
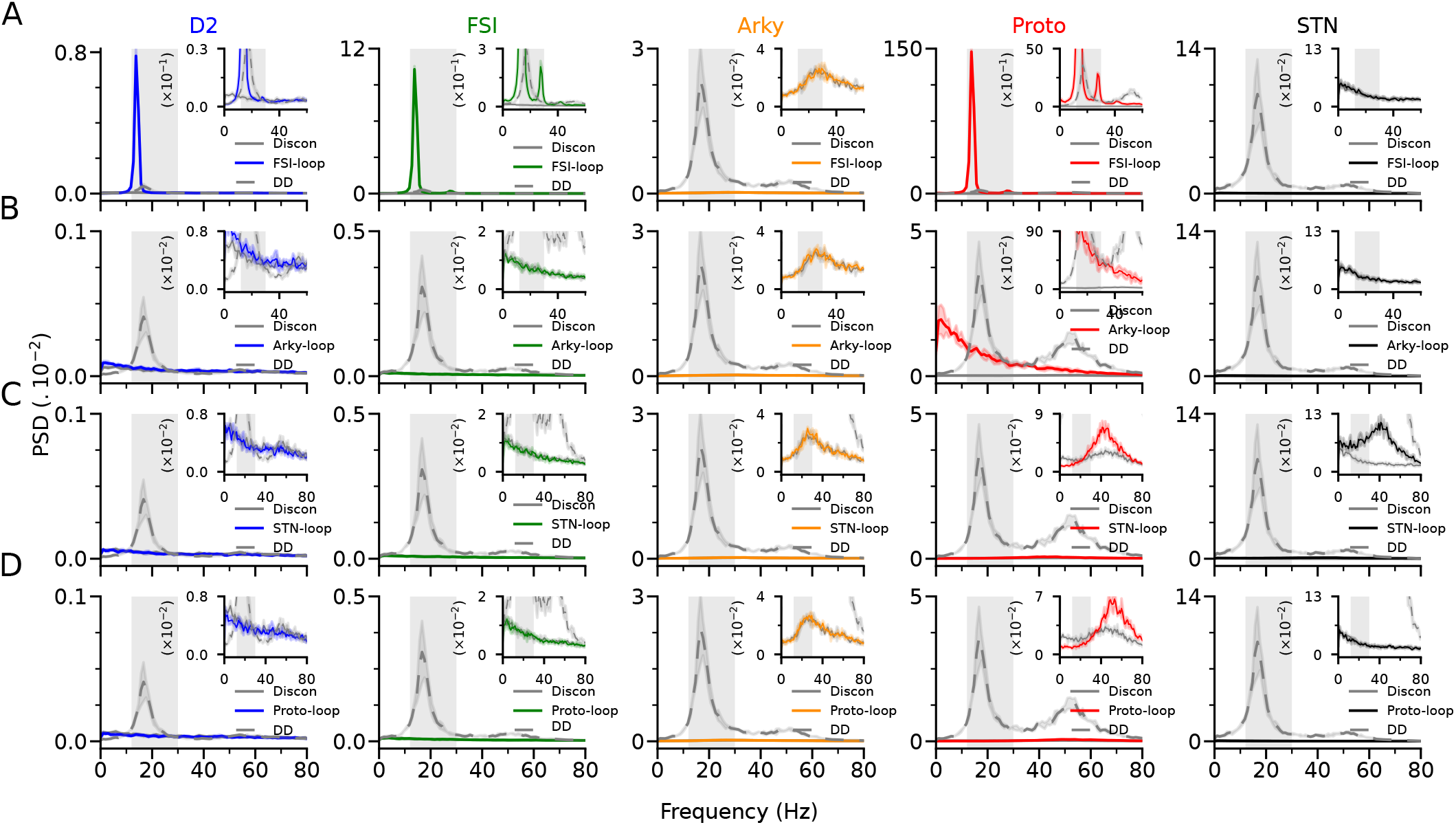
*β* oscillations result from synaptic changes due to DD in the FSI-D2-Proto loop that pushes it to oscillatory regime. A) The power spectrum densities of the population firing rate in different states. Solid colored lines correspond to network activity with only the FSI-D2-Proto loop included. Whereas the grey dash-dotted line represents the DD state with all the loops included (as in Fig. 7C). The solid grey line represents a network with no synaptic connection i.e. ”disconnected”. The insets show the same results in higher resolution. B) Same as in (A) but including only the Arky-D2-Proto loop. C) Same as in (A) but for the STN-GPe(Proto) loop. D) Same as in (A) but for the Proto-Proto loop.

